# Lineage-dependence of the neuroblastoma surfaceome defines tumor cell state-dependent and independent immunotherapeutic targets

**DOI:** 10.1101/2024.06.27.600865

**Authors:** Nathan M. Kendsersky, Michal Odrobina, Nathaniel W. Mabe, Alvin Farrel, Liron Grossmann, Matthew Tsang, David Groff, Adam J. Wolpaw, Francesca Zammarchi, Patrick H. van Berkel, Chi V. Dang, Yaël P. Mossé, Kimberly Stegmaier, John M. Maris

**Affiliations:** Division of Oncology and Center for Childhood Cancer Research; Children’s Hospital of Philadelphia; Philadelphia, PA, 19104; USA; Perelman School of Medicine at the University of Pennsylvania; Philadelphia, PA, 19104; USA; Department of Pediatric Oncology; Dana Farber Cancer Institute; Boston, MA, 02215; USA; Department of Biomedical and Health Informatics (DBHi); Children’s Hospital of Philadelphia; Philadelphia, PA, 19104; USA; ADC Therapeutics (UK) Ltd, London; United Kingdom; Department of Oncology, Johns Hopkins University School of Medicine; Department of Biochemistry and Molecular Biology, Johns Hopkins Bloomberg School of Public Health; Baltimore, MD, 21218; USA; Ludwig Institute for Cancer Research, New York, NY 10016; USA; The Broad Institute of MIT and Harvard; Cambridge, MA, 02142; USA

**Keywords:** neuroblastoma, immunotherapy, epigenetics, surfaceome, AXL, antibody-drug conjugate

## Abstract

**Background:** Neuroblastoma is a heterogeneous disease with adrenergic (ADRN)- and therapy resistant mesenchymal (MES)-like cells driven by distinct transcription factor networks. Here, we investigate the expression of immunotherapeutic targets in each neuroblastoma subtype and propose pan-neuroblastoma and cell state specific targetable cell-surface proteins.

**Methods:** We characterized cell lines, patient-derived xenografts, and patient samples as ADRN-dominant or MES- dominant to define subtype-specific and pan-neuroblastoma gene sets. Targets were validated with ChIP-sequencing, immunoblotting, and flow cytometry in neuroblastoma cell lines and isogenic ADRN-to-MES transition cell line models. Finally, we evaluated the activity of MES-specific agents *in vivo* and *in vitro*.

**Results:** Most immunotherapeutic targets being developed for neuroblastoma showed significantly higher expression in the ADRN subtype with limited expression in MES-like tumor cells. In contrast, *CD276* (B7-H3) and *L1CAM* maintained expression across both ADRN and MES states. We identified several receptor tyrosine kinases (RTKs) enriched in MES-dominant samples and showed that AXL targeting with ADCT-601 was potently cytotoxic in MES-dominant cell lines and showed specific anti-tumor activity in a MES cell line-derived xenograft.

**Conclusions:** Immunotherapeutic strategies for neuroblastoma must address the potential of epigenetic downregulation of antigen density as a mechanism for immune evasion. We identified several RTKs as candidate MES-specific immunotherapeutic target proteins for the elimination of therapy-resistant cells. We hypothesize that the phenomena of immune escape will be less likely when targeting pan-neuroblastoma cell surface proteins such as B7-H3 and L1CAM, and/or dual targeting strategies that consider both the ADRN- and MES-cell states.

**Key Points:**

- Cellular plasticity influences the abundance of immunotherapeutic targets.
- Subtype-specific targets may be susceptible to epigenetically-mediated downregulation.
- Immunotherapeutic targets in development, B7-H3 and L1CAM, show “pan-subtype” expression.

**Importance of Study:** Neuroblastoma is a lethal childhood malignancy that shows cellular plasticity in response to anti-cancer therapies. Several plasma membrane proteins are being developed as immunotherapeutic targets in this disease. Here we define which cell surface proteins are susceptible to epigenetically regulated downregulation during an adrenergic to mesenchymal cell state switch and propose immunotherapeutic strategies to anticipate and circumvent acquired immunotherapeutic resistance.

## Introduction

Neuroblastoma is a pediatric extracranial solid tumor that arises from deregulated development of the sympathoadrenal nervous system.^1,2^ It is characterized by clinical heterogeneity ranging from low-risk tumors that can spontaneously regress to extremely aggressive high-risk tumors that relentlessly progress despite intensive multimodal therapy. Neuroblastoma is currently the only pediatric solid tumor with a labeled immunotherapy indication for monoclonal antibodies specific to the GD2 glycolipid abundantly present on the surface of neuroblastoma cells.^3–6^ These monoclonal antibodies, however, show significant on-target, off-tumor toxicities as GD2 is expressed on pain fibers and this glycolipid can be epigenetically downregulated as a potential mechanism of resistance.^7^ Chimeric antigen receptor (CAR) engineered autologous T cell therapies targeting GD2 have shown recent impressive anti-tumor activity in neuroblastoma,^8^ and diffuse midline gliomas,^9^ however in both responses are often transient.

Despite intensive chemotherapy, radiation therapy, and immunotherapy, approximately 50% of patients with high-risk neuroblastoma suffer relapse which remains generally incurable, highlighting the need to identify new therapeutic targets. Investigators have evaluated the activity small molecule inhibitors, radiotherapies, and immunotherapeutic agents targeting neuroblastoma-specific molecular aberrations. Lorlatinib, an inhibitor of mutationally activated anaplastic lymphoma kinase (*ALK*), has shown potent preclinical and clinical activity,^10–12^ and is now being studied in a Phase 3 trial for newly diagnosed high-risk patients harboring an *ALK* mutation (NCT031226916). ALK is also the target of various immunotherapeutic strategies in preclinical development, including CARs,^13,14^ and antibody drug conjugates (ADCs).^15^ The norepinephrine transporter (NET, encoded by *SLC6A2*) is another protein of interest that is used for both imaging and therapeutic purposes.^2^ Radiolabeled ^131^I-MIBG, which is shuttled into neuroblastoma tumor cells by the norepinephrine transporter overexpressed on the cell surface, is also being studied in the same Phase 3 trial as lorlatinib. *L1CAM* (CD171) is the gene encoding a cellular adhesion molecule overexpressed in neuroblastoma and is the target of a recent CAR T trial.^16,17^ *CD276* encodes B7-H3, which is another cell-surface molecule overexpressed in a variety of pediatric solid tumors. Monoclonal antibodies, ADCs, and CARs specific to B7-H3 are being explored in preclinical and clinical studies.^18–21^ GPC2 is a glypican protein anchored to the cell surface that was discovered as an oncogene in neuroblastoma and has been targeted with ADC and a CAR T cell approaches^22–25^, including an ongoing clinical trial (NCT05650749). It is also important to understand how heterogeneity and plasticity affect expression of immunotherapeutic targets such that we can prioritize the many therapeutic strategies in development and to uncover novel targets enriched in therapy-resistant cells.

Neuroblastoma cells can exist as genetically identical but epigenetically distinct states, as originally described decades ago.^26,27^ Cells can be categorized as adrenergic (ADRN) or mesenchymal (MES) as defined by distinct epigenetically regulated core regulatory circuit (CRC) transcription factors that can transdifferentiate in cell line models.^28–33^ Single-cell studies of developing neural crest cells in mice have described these two developmental branches and their associated lineage-related transcription factors.^34^ The clinical relevance of this plasticity is postulated to be in therapy resistance as was recently demonstrated with downregulation of GD2 in response to GD2-directed immunotherapies.^7^ Here, we sought to define the plasticity of the neuroblastoma surfaceome in light of several emerging immunotherapeutic strategies for this disease.

## Materials and Methods

### RNA-sequencing Data

Neuroblastoma RNA sequencing data was accessed through the Treehouse Childhood Cancer Initiative and Gabriella Miller Kids First (GMKF) Pediatric Research Program. Treehouse data was obtained as expected count matrices aligned to Hg38 (Treehouse Tumor Compendium v11 Public PolyA https://treehousegenomics.soe.ucsc.edu/public-data/#tumor_v11_polyA, N = 200 tumors). GMKF data was collected from the GMKF Data Resource Center (https://d3b.center/kidsfirst/, N = 195 samples). We only assessed samples that were confirmed as neuroblastoma by histopathology. Thirteen patient samples were represented in both the Treehouse and GMKF datasets. For differential expression analysis, Treehouse and GTEx datasets were converted to counts per million (cpm), filtered by genes with cpm>1, and normalized with the Empirical Analysis of Digital Gene Expression Data package (edgeR) (trimmed mean of M-values, or TMM) for downstream analyses.^35,36^ We also utilized our RNA-sequencing data for parental neuroblastoma cell lines (N=38) and patient-derived xenografts (N=30) as previously described.^37,38^ RNA-sequencing for ADRN-to-MES transition models (SKNBE2 and KPNYN) was obtained from [GSE180516].^7^

### Subtype Classification

Neuroblastoma samples (Treehouse human tumors, GMKF human tumors, and neuroblastoma cell lines) were analyzed with singscore, a single sample gene set enrichment analysis (ssGSEA) method.^39^ The gene set lists for ADRN and MES cell states were obtained from Groningen and colleagues.^29^

### Differential Expression (limma)

Log2TPM RNA-seq data were transformed with the *voom* function and modeled with lmfit.^40^ Contrasts were prepared between each neuroblastoma subtype. For each comparison, the linear model was evaluated with limma using empirical Bayes Statistics (eBayes) and adjustment with Benjamin-Hochberg post hoc testing.^41^ To evaluate ADRN-specific and MES-specific differentially expressed genes, we filtered by a minimum Log2 fold change >1 and max adjusted p value <0.05. Pan-subtype genes were defined by a minimum Log2 fold change <1 and max adjusted p value >0.05. Surface proteins were annotated with the UniProt localization database with the keyword “Cell Membrane”. Subtype-specific and pan-subtype genes were also intersected with the Food and

Drug Administration’s Pediatric Relevant Molecular Target List for childhood cancers (https://www.fda.gov/media/120332/download).^42^

### Deconvolution

Two algorithms were used to predict cellular fractions of bulk patient RNA-sequencing data. quanTIseq was performed with bulk TPM data and the --tumor=TRUE flag.^43^ EPIC was performed with bulk TPM data after removing common genes in both MES-specific neuroblastoma and cancer-associated fibroblast signatures (COL1A1, COL3A1, SYNPO2).^44^

### ChIP-sequencing Analysis

ChIP-sequencing data for H3K4me1, H3K4me3, H3K27Ac, and H3K27me3 were obtained from the GEO database (GSE138315).^45^.

### DNA-methylation Analysis

Methyl-sequencing data were obtained from the TARGET project available on the NCI Office of Cancer Genomics website (https://target-data.nci.nih.gov/Public/NBL/methylation_array/).^46^ Samples with available mRNA-sequencing data were used to determine dominant subtype (120 ADRN samples and 8 MES samples). The R package gviz was used to generate figures.

#### GSEA

The *fgsea* and *msigdbr* packages were utilized to find gene sets associated with both ADRN and MES Treehouse and GMKF patient tumors.^47,48^

### Cell Culture

Neuroblastoma cell lines were cultured in RPMI 1640 Medium with L-glutamine (Corning cat. 10041CV) with 10% FBS (VWR cat. 89510-186) and 1% L-glutamine (Corning cat. 25005CI) or IMDM (ThermoFisher cat. 12440053) with 20% FBS, 1% L-glut, and 1:1000 ITS+ Premix Universal Culture Supplement (Corning cat.

354352). When culturing cell lines with doxycycline-regulated transgenes, we used Tet-Free FBS (Takara cat. 631106).

For CRISPR/Cas9 knockout of *AXL*, we utilized the lentiCRISPRv2 vector (Addgene cat. 52961) with one of two *AXL*-targeted sgRNAs (Oligo 1: CACCGCTGAGAACATTAGTGCTACG; Oligo 2: CACCGGCTGCTGGTGCATGCCACG) ^49^. We also cloned a non-targeting (sgLacz: CACCGAACGGCGGATTGACCGTAAT) and an off target sgRNAs (Chr2.2: CACCGGTGTGCG-TATGAAGCAGTG). lentiCRISPRv2 constructs were independently packaged into lentiviral particles in HEK293T cells with psPAX2 (Addgene cat. 12260) and pMD2.G (Addgene cat. 12259), according to the Lipofectamine 2000 protocol (Invitrogen cat. 11668027). Cell line models were transduced with lentiviral particles and polybrene (10µg/mL) containing one of four lentCRISPRv2 constructs and selected with puromycin (1µg/mL) for several weeks.

AXL overexpression was achieved with the TetR protein and a CMV/TO-regulated *AXL* transgene. We used Gateway Clonase II (Invitrogen cat. 11791020) to insert the *AXL* cDNA (GeneCopoeia cat. GC-Z7835) into pLenti-CMV/TO-puroR-DEST (Addgene cat. 17293). To regulate expression of the CMV/TO-AXL transcript, we utilized the pLenti-CMV-TetR-blastR construct (Addgene cat. 17492). Both pLenti-CMV/TO-AXL-puroR and pLenti-CMV-TetR-blastR constructs were independently packaged into lentiviral particles, as above. Parental neuroblastoma cell lines were successively transduced with the TetR-containing lentivirus, selected with 5µg/mL blasticidin (Invitrogen cat. R21001), transduced with the CMV/TO-AXL lentivirus, and finally selected with 1µg/mL puromcyin (Gibco cat. A1113803). Expression of *AXL* was induced by adding 1µg/mL of doxycycline (Sigma cat. D3072) to the culture medium.

### Sample Processing and Immunoblotting

Whole-cell lysates were prepared using RIPA Lysis Buffer System (Santa Cruz Biotechnology cat. sc-24948), Protease Inhibitor Cocktail (Sigma-Aldrich cat. P8340), and Phosphatase Inhibitor Cocktails 2 and 3 (Sigma-Aldrich cat. P5726 and cat. P0044). Pellets were resuspended in lysis buffer, incubated on ice for 15 minutes, disturbed with a vortex for 20 seconds, and then cleared by centrifugation at max speed (17900 x g) for 15 minutes at 4°C. Protein concentration was determined according to the Bio-Rad Protein Assay Kit II (Bio-Rad cat. 5000006). Once quantified, 15µg of each protein sample was prepared with Laemmli Sample Buffer (Bio-Rad cat. 1610737) and 50mM DTT (Millipore Sigma cat. 43816). Samples were loaded on a 4-15% Criterion TGX Protein Gel (Bio-Rad cat. 5671085) and run with Tris-Glycine Buffer (Bio-Rad cat. 1610771) containing 0.1% SDS (Invitrogen cat. 15553027) at 40mAmps per gel. Proteins were transferred to a 0.45µm Immobilon-P Membrane (Millipore Sigma cat. IPVH00010) in ice-cold Tris-Glycine Buffer with 20% methanol in a transfer tank either overnight at 10V or 1 hour at 50V. Membranes were blocked with 5% Blotting-Grade Blocker (Bio-Rad cat. 1706404) in Tris Buffered Saline with Tween® 20 (TBST, Cell Signaling Technology cat. 9997) for 1 hour and incubated with primary antibody in 5% Blocking Buffer overnight at 4°C. Primary Antibodies and dilutions are as follows: ALK (1:1000, CST cat. 3333), DLL3 (1:1000, CST cat. 78110), GPC2 (1:500, SCBT cat. Sc- 393824), SLC6A2 (1:500, MAb Technologies cat. NET17-1), CD276 (1:1000, Abcam cat. ab134161), L1CAM (1:1000, CST cat. 89861), AXL (1:200, R&D Systems cat. AF154), EGFR (1:1000, Thermo Fisher cat. MA5- 13070), EPHA2 (1:1000, Thermo Fisher cat. 37-4400), PDGFRA (1:1000, Thermo Fisher cat. 710169), PDGFRB (1:1000, Thermo Fisher cat. MA5-15143), PHOX2A (1:1000, SCBT cat. sc-81978), DBH (1:1000, CST cat. 8586), SNAI2 (1:1000, CST cat. 9585), VIM (1:1000, CST cat. 5741). Membranes were then washed 4 times for 10 minutes with TBST. We then incubated membranes with secondary antibody (in 5% Blocking Buffer) for 1 hour at room temperature (Invitrogen cat. A16110; Invitrogen cat. 31432; Invitrogen cat. 31402; R&D Biosystems cat. HAF016). We performed a final TBST wash (4 x 10 minutes) before incubating membranes in Pierce ECL Plus (Thermo Scientific cat. 32132) or SuperSignal West Femto ECL (Thermo Scientific cat. 34095) for 5 seconds to 3 minutes. We imaged membranes using the Azure Biosystems Sapphire^TM^ Biomolecular Imager.

### Flow Cytometry

Neuroblastoma cell lines were counted and distributed into FACS tubes for various staining conditions and an unstained control (0.5M cells / condition). Cells were washed 3 times with 1mL of PBS, spinning at 300 x g for 5 minutes at 4°C between each step. Staining solutions were prepared in PBS with the manufacturer’s recommended antibody and LIVE/DEAD Fixable stain (Invitrogen cat. L34980) amounts per assay. Conjugated primary antibodies include the following: APC-AXL (R&D Systems cat. FAB154A), PE-EphA2 (BioLegend cat. 356803), APC-PDGFRA (BioLegend cat. 323511), AF488-HER2 (BioLegend cat. 324410), BV421-EGFR

(BioLegend cat. 352911), BV605-PDGFRB (BD Biosciences cat. 743035). Cells were resuspended in 100µL of the appropriate staining buffer and incubated on ice and in the dark for 30 minutes. After the antibody incubation, cells were again washed 3 times with 1mL of PBS, spinning at 300 x g for 5 minutes at 4°C between each step. Cells were fixed in 1% formaldehyde ice and in the dark for 15-30 minutes, then washed 2 times with 1mL of PBS. Finally, cells were resuspended in 100µL of PBS and stored at 4°C in the dark until samples were analyzed on a CytoFLEX LX Flow Cytometer (Beckman Coulter). Data was analyzed using FlowJo® software.

### Cytotoxicity Assays

Small molecule inhibitors were obtained from Selleck Chemicals: Bemcentinib (S2841), Cabozantinib (S1119), NPS-1034 (S7669), ONO-7475 (S8933), Erlotinib (S7786), Crenolanib (S2730), Afatinib (S1011), Imatinib (S2475), Sitravatinib (S8573), Dasatinib (S1021), Erdafitinib (S8401), ALW II-41-27 (S6515), Sapitinib (S2192). AXL-targeting (ADCT-601) and control (B12-PL1601) antibody-drug conjugates (ADCs) were studied in collaboration with ADC Therapeutics ^50^. Neuroblastoma cell lines were seeded in a 96-well plate one day prior to treatment with small molecule inhibitors or antibody-drug conjugates. Assays with small molecule inhibitors (128pM – 50µM) were concluded after 96 hours, while ADCs studies with (0.667fM – 66.7nM) were evaluated after 120 hours. At the study endpoint, cellular viability was determined using the CellTiter-Glo 2.0 Assay protocol (Promega cat. G9243). Luminescence values were normalized to untreated wells and data was analyzed using R 4.0.3 (2020-10-10). Plots and IC50 values were calculated using the following packges: ggplot2 (v3.3.5), nplr (v0.1-7), drc (v3.0-1), dr4pl (v2.0.0), tidyverse (v1.3.1). Each experiment was plated in technical triplicate and data are representative of at least two independent experiments.

### In Vivo Studies

For murine efficacy studies, we engrafted CB17 severe combined immunodeficiency (SCID) mice with cell line-derived xenografts (CDXs) or patient-derived xenografts (PDXs) and followed the standard protocol established in the Pediatric Preclinical Testing Consortium (now named Pediatric Preclinical In Vivo Testing, PIVOT).^19^ In brief, mice with tumors at enrollment size (0.2-0.3 cm^3^) received 1 mg/kg of either ADCT-601 or B12-PL1601 via tail vein injection. Studies were performed with N=6 animals per arm and the mice were monitored for 100 days or until their tumor burden reached 2.0 cm^3^.

## Results

*Human neuroblastoma tumors and cell lines classified as ADRN- or MES-dominant display differentially expressed genes beyond previously established gene signatures.* Prior studies defined a 485-gene MES signature and a 369-gene ADRN signature and these have been widely used to characterize these states.^29^ However, these signatures were derived from a relatively small number of cell lines. We asked if we could use these signatures to identify ADRN and MES tumors and cell lines in larger datasets, then analyze those datasets to identify additional genes expressed in a state-specific manner. We assessed two datasets including the predominantly high-risk samples in the Treehouse Childhood Cancer Initiative (Treehouse) and slightly non-high-risk tumor biased case series in the Gabriella Miller Kids First dataset (**Supplemental Figure 1a**). Using single sample gene set enrichment analysis (ssGSEA, singscore), we evaluated the ADRN and MES score for each tumor and characterized each sample by the dominant signature. (**Figure 1a, Supplemental Figure 1b**). Most tumors in the Treehouse and GMKF databases were predominantly ADRN (88.4% and 92.8% respectively), though several tumors in each dataset were MES-dominant. Within each patient dataset, risk group was not appreciably different between ADRN and MES subsets (**Supplemental Figure 1c**). We also determined the dominant subtype in a neuroblastoma cell line dataset,^37^ and the neuroblastoma models in the Cancer Cell Line Encyclopedia (CCLE),^51^ finding that 94.7% and 83.3% were classified as ADRN, respectively (**Supplemental Figure 1d-e**).

**Figure 1.**
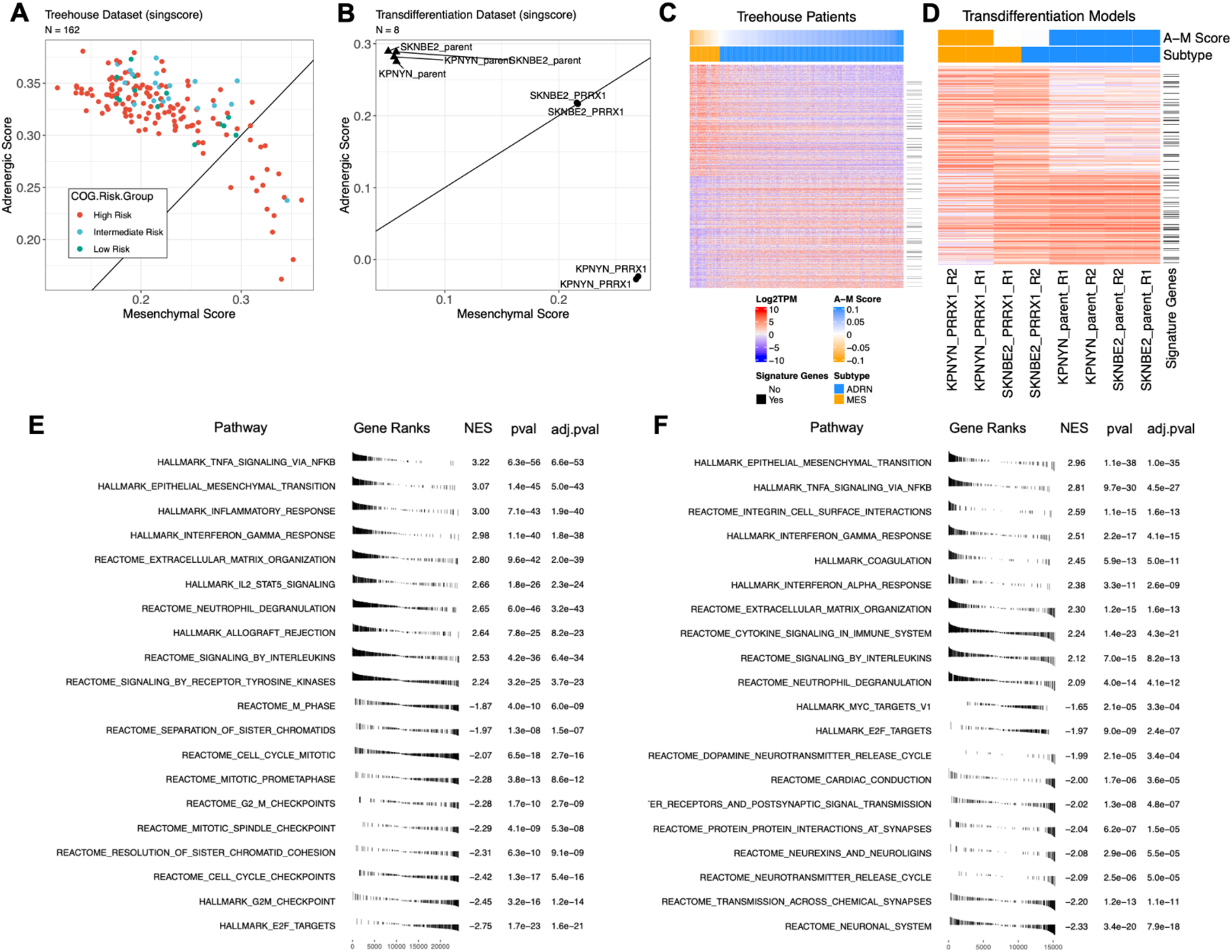
A-B) Single-sample gene set enrichment analysis and published ADRN/MES gene signatures were used to generate ADRN and MES scores to classify (A) Treehouse patient samples and (B) ADRN-to-MES transdifferentiation models as ADRN- or MES-dominant. Patient samples are colored by COG Risk Group. Transition models represent two replicates of each model (SKNBE2 and KPNYN; parent and PRRX1). C) Heatmap displaying the top 500 MES and top 500 ADRN differentially expressed genes, as calculated by limma/voom. D) Heatmap displaying data from transdifferentiation models, filtered for differentially expressed genes from the Treehouse patient dataset. ADRN/MES signature genes are denoted by a horizontal black line. A-M (ADRN minus MES) scores and subtypes were defined by singscore. E-F) Gene set enrichment analysis (fgsea) of MES- and ADRN-dominant (E) Treehouse patient samples and (F) ADRN-to-MES transdifferentiation models using Hallmark and curated (C2) MSigDB pathways. Values represent normalized enrichment scores (NES), p-values (pval, based on multi- level split Monte-Carlo scheme), and Benjamini-Hochberg-adjusted p-values (adj.pval).

We also evaluated top ADRN and MES differentially expressed genes in an ADRN-to-MES transdifferentiation (or transition) model, whereby inducible overexpression of the *PRRX1* MES transcription factor can convert ADRN cell lines to a MES-like phenotype.^7^ By singscore, both SKNBE2 and KPNYN parental cell lines assayed in this system were ADRN, with average ADRN and MES scores of 0.243 and 0.0552, respectively (**Figure 1b**). Upon overexpression of the *PRRX1* transgene, SKNBE2 cell lines showed a slightly decreased mean ADRN score (0.217) and significantly increased mean MES score (0.216), classifying SKNBE2- PRRX1 as a mixed phenotypic model. KPNYN cell lines overexpressing PRRX1 transitioned completely from ADRN to MES, with average ADRN and MES scores of -0.0253 and 0.269, respectively.

To further characterize neuroblastoma tumor subtypes, we performed differential gene expression between ADRN- and MES-predominant neuroblastoma Treehouse samples with limma/voom methodologies. We observed 4380 MES and 4199 ADRN differentially expressed genes (DEGs) using a filtering strategy of log2 fold change >1 and adjusted p value <0.05. The MES DEGs included 320 of 485 (83.1%) MES signature genes and the ADRN DEGs included 153 of 369 (41.5%) ADRN signature genes. Of the top 1000 differentially expressed genes between ADRN- and MES-dominant Treehouse samples, only 75 were present in the published ADRN/MES gene signatures (**Figure 1c**), suggesting there are more features defining these distinct phenotypes. We also show that the same ADRN and MES DEGs in Treehouse samples were differentially expressed in the GMKF and cell line datasets (**Supplemental Figure 1f**). The ADRN and MES DEGs obtained from the patient dataset are also reflected in the transdifferentiation models (**Figure 1d)**.

We next sought to validate these findings with histone ChIP-sequencing data because neuroblastoma subtypes are epigenetically mediated. We evaluated the loci of differentially expressed genes for the presence of active enhancer/promoter (H3K27ac) and repressive (H3K27me3) histone marks (**Supplemental Figure 1g**). We showed that enhancer marks bound to subtype-specific differentially expressed genes, where H3K27ac marks are preferentially deposited at ADRN loci in ADRN models and MES loci of the MES model. As expected, ADRN DEGs are bound by repressive marks in the MES cell line model, while the loci of MES DEGs are repressed in ADRN models. Altogether, this suggests that these patient samples and cell line models display similar transcriptional and epigenetic states, which can help us better understand the ADRN and MES phenotypes as they relate to relevant therapeutic approaches.

As the majority of differentially expressed genes were not in the published ADRN/MES gene signature,^29^ we performed Gene Set Enrichment Analysis to identify pathways associated with neuroblastoma patient tumors and transition cell line models (**Figure 1e-f**). We assessed the Hallmark gene sets and C2 curated gene sets (including BIOCARTA, KEGG, REACTOME) from the Molecular Signatures Database (MSigDB). In both MES- dominant patient samples and transition models, the top gene sets were related to immune cell signaling, including TNF-α, IFN-γ, and inflammatory response (**Figure 1e-f**), which validated recently published data^52,53^. ADRN-dominant patient samples were enriched cell cycle-related gene sets including Hallmark pathways for E2F targets and G2/M checkpoints, as well as Reactome pathways for cell cycle checkpoints, G2/M checkpoints, M-phase, and others (**Figure 1e-f**).

We performed deconvolution with quanTIseq and EPIC to evaluate differences in immune cell subsets between MES and ADRN tumors. While some of our findings were confounded by the presence of immune cell signatures in cell lines (**Supplementary** Figure 2a), our analysis using EPIC showed ADRN-dominant tumors were composed of more tumor cells (7.75e-07) and MES-dominant tumors were enriched in macrophages (2.69e-03) and cancer associated fibroblasts (CAFs, p = 2.32e-06) (**Supplemental Figure 2b**). The finding of MES-enriched CAFs supports previous publications describing an association between M2 tumor-associated macrophages and CAFs, which increased the chemotherapeutic resistance of neuroblastoma tumor cells *in vitro.*^54^

Next, we sought to validate the MES-associated immune-related pathways *in vitro*. In addition to B2M and HLA-A/B/C previously identified in the MES signature,^29^ we showed MES-specific expression of additional antigen processing and presentation genes (*HLA-E* and *TAPBP*), the immunomodulatory protein PD-L1 (*CD274*), and cytokines CCL2 and CSF1 (**Figure 2a-b**). This pattern was also observed at the epigenomic level, as *HLA-A*, *HLA-B*, *HLA-C*, *HLA-E*, *CD274*, and *CCL2* were bound by H3K27ac markers only in the SKNAS mesenchymal model (**Supplemental Figure 3**). Conversely, ADRN cell lines were bound by repressive histone marks at these same loci. We validated cell surface expression of MHC Class I and PD-L1 in MES-like parental cell lines and the ADRN-to-MES transition models (**Figure 2c-d**), as well as increased CCL2 secretion in MES parental and transition cell line models (**Figure 2e**). Together, these data suggest that pathways critical for adaptive immunity are suppressed in the ADRN cell state and are upregulated in MES-like neuroblastoma cells.

**Figure 2.**
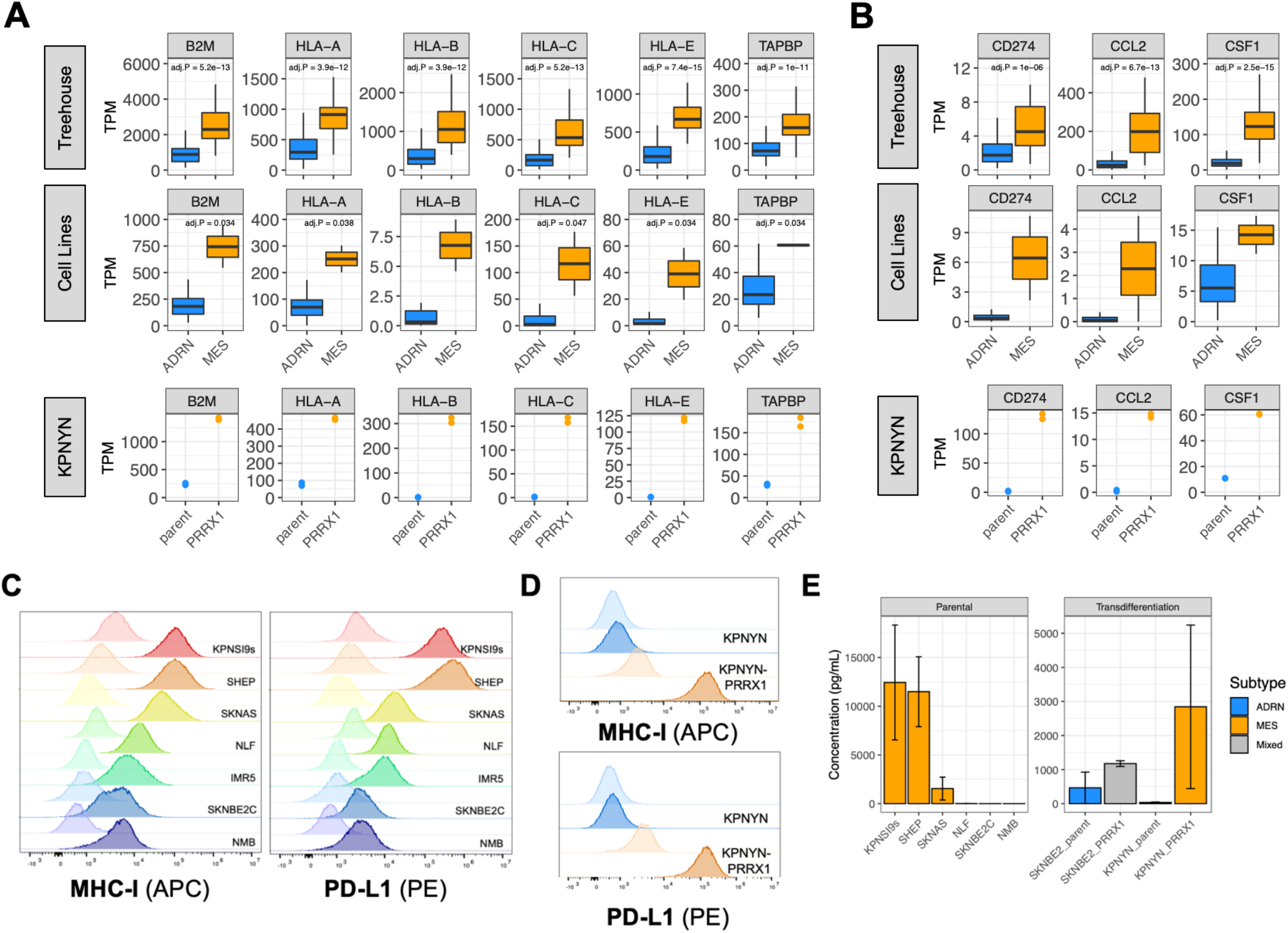
Immune signatures, including antigen presentation pathway and immunomodulatory cytokines, are enriched in MES-like neuroblastoma samples. A-B) RNA expression of (A) antigen presentation pathway genes in Treehouse Patient samples, cell lines, and the KPNYN cell line before (parent) and after (PRRX1) transdifferentiation. C-D) Cell surface staining of MHC-I and PD-L1 in (C) parental and (D) ADRN-to-MES transition models. Lighter colors represent unstained control. (E) Quantification of extracellular CCL2 in (left) parental and (right) transdifferentiation cell line models (N = 2, error bars show standard error of the mean).

*Most known cell-surface therapeutic targets are differentially expressed between neuroblastoma ADRN and MES subtypes.* In the ADRN-dominant patient samples and cell lines, we observed subtype-specific RNA expression of candidate immunotherapeutic targets, including *ALK, GPC2*, and *DLL3,* as well as the targeted radiotherapeutic target *SLC6A2* (NET) (**Figure 3a, Supplemental Figure 4a-b**), including subtype-specific deposition of enhancer histone marks at each gene locus (**Supplemental Figure 4c**). We did not observe differential DNA methylation patterns at select subtype-specific loci between ADRN (N=120) and MES (N=8) samples (**Supplemental Figure 4d-e**). We also observed ADRN-specific expression of ALK at the protein level (**Figure 3b**). We validated these findings by showing that ADRN-to-MES transdifferentiation results in a loss of ADRN-specific target gene expression of these therapeutically tractable proteins (**Figure 3a,c** **and Supplemental Figure 4a-b**). In the KPNYN model with a complete subtype switch to MES, expression of *ALK*, *DLL3*, *GPC2*, and *SLC6A2* were substantially decreased to complete or near complete loss. In the SKNBE2 cell line with gain of MES signature and retention of ADRN signature, the expression of these therapeutic target genes decreased partially or did not change. Overall, these data suggest that these clinical and preclinical cell surface therapeutic targets are enriched in ADRN-predominant cells and are downregulated in MES-predominant cells.

**Figure 3.**
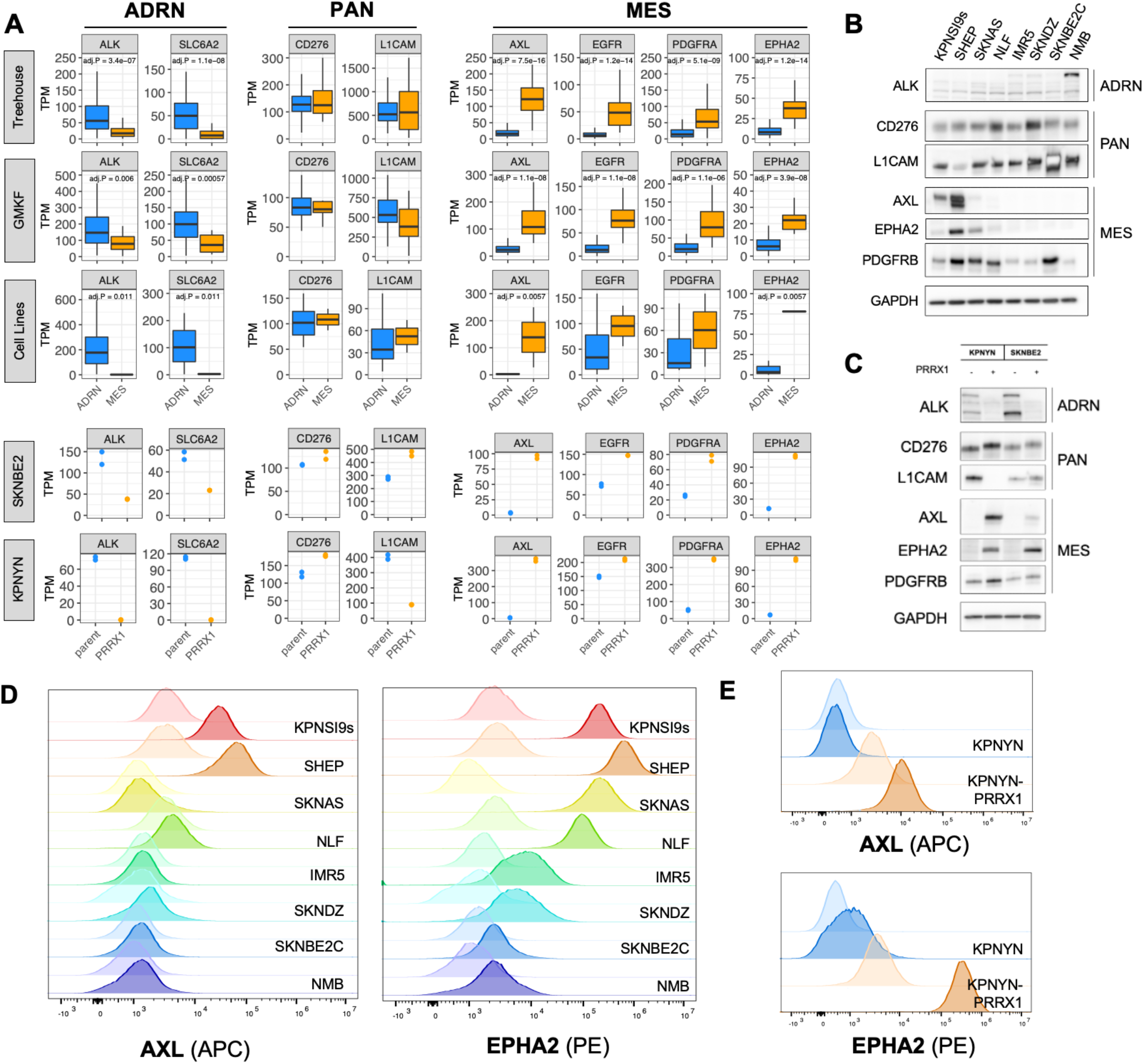
Differential expression reveals ADRN-specific, pan-subtype, and MES-specific targets. A) RNA expression of ADRN-specific, pan-subtype, and MES-specific targets in patient tumors, parental cell lines, and ADRN-to-MES transition models (SKNBE2 and KELLY). B-C) Protein expression of select targets in (B) parental and (C) ADRN-to-MES transdifferentiation cell line models. D-E) Cell surface staining of AXL and EPHA2 in (D) parental and (E) ADRN-to-MES transition cell line models. Flow cytometry figures are comparing unstained (lighter) to stained (darker) for each sample.

In contrast, *CD276* and *L1CAM* were similarly expressed across both MES and ADRN subtypes (**Figure 3a**). We supported these findings by showing pan-subtype enhancer marks and protein expression across a panel of neuroblastoma cell line models. (**Figure 3b**, **Supplemental Figure 4c**). Further, *CD276* mRNA and protein expression increased or stayed the same in both KPNYN and SKNBE2 transdifferentiation models assayed, while *L1CAM* showed discordant results (**Figure 3a,c**). Overall, these data support B7-H3 and L1CAM as subtype independent neuroblastoma immunotherapeutic targets.

We assessed MES-specific genes that were present in the Pediatric Relevant Molecular Target List to enrich for proteins that are recognized as potential therapeutic targets across childhood cancers.^42,55^ We showed that the receptor tyrosine kinases *AXL*, *EGFR*, *PDGFRA*, and *EPHA2* were specifically enriched in MES-dominant patient samples, as well as MES neuroblastoma cell lines (**Figure 3a**). We observed H3K27ac enhancer marks at the loci of these MES-specific receptor tyrosine kinases (RTKs) only in the MES neuroblastoma cell lines, while ADRN models displayed H3K27me3 marks (**Supplemental Figure 4c**). This effect was particularly pronounced at the *AXL* and *EPHA2* loci. At the protein level, AXL, EPHA2, and PDGFRB showed MES-specific expression (**Figure 3d**). Despite some variation at the level of histone marks, transition from ADRN-to-MES resulted in significant upregulation of MES-specific RTK transcripts (**Figure 3a**), as well as protein measured by immunoblotting and flow cytometry (**Figures 3c,e**).

*MES-specific RTKs are therapeutic vulnerabilities in MES neuroblastoma cell line models.* AXL is linked to the EMT gene signature, and by altering the cell signaling pathways, AXL and other RTKs may confer resistance to chemotherapy and targeted therapies.^56–58^ We sought to further understand AXL and its expression on tumor and stromal cells with our deconvolution algorithm. As AXL was minimally expressed in the ADRN subtype, we did not observe any correlations between AXL expression and tumor/immune-cell fraction in ADRN-dominant samples (**Supplemental Figure 5**). We did observe a negative correlation between AXL expression and tumor fraction in MES-dominant samples (cor = -0.5; p = 6.85e-3) (**Supplemental Figure 5a**), though the MES- dominant samples with the highest tumor purity still expressed AXL at higher levels than ADRN-dominant samples. We showed that AXL expression is positively correlated with monocytes in the MES-dominant tumor samples (cor = 0.48; p = 1.04e-2; **Supplemental Figure 5b**). Together, these data and our *in vitro* data (**Figure 3**) suggest that AXL is present on both neuroblastoma cells and cell populations within the tumor microenvironment.

To assess the functional role of AXL on tumor cells, we modulated its expression in neuroblastoma cell line models and monitored the impact on subtype-specific markers.After doxycycline-inducible *AXL* overexpression in neuroblastoma cell lines, we observed subtle changes in the IMR5 model whereby some MES markers slightly increased and some ADRN markers slightly decreased (**Supplemental Figure 6a-c**), supporting AXL as a contributor to the MES subtype. After complete *AXL* knockout using CRISPR/Cas9, we did not observe changes in MES or ADRN markers (**Supplemental Figure 6d-f**). These data suggest that AXL is not solely necessary for maintenance of the MES subtype, but likely contributes in combination with other signaling proteins.

We next evaluated the activity of several MES-specific RTK small molecule inhibitors in development. For AXL, we examined the cytotoxicity of bemcentinib, cabozantinib, NPS-1034, and ONO-7475 in a panel of ADRN and MES neuroblastoma cell lines. While bemcentinib was similarly cytotoxic in both ADRN and MES models, the remaining AXL inhibitors were slightly more active in MES models (**Figure 4a**). Despite increased cytotoxicity in MES models, the drugs were not especially potent, as the minimum IC50 values for cabozantinib, NPS-1034, and ONO-7475 were 1.1µM, 876nM, and 144nM respectively (**Figure 4c**). Additionally, depletion of AXL did not result in resistance to AXL inhibition (**Supplemental Figure 7**), suggesting that cytotoxicity was mediated in part by nonspecific inhibition of other receptor tyrosine kinases. The EPHA2 inhibitor ALW II-41-27 and the multi-target inhibitor dasatinib showed MES-specific activity across a panel of parental neuroblastoma cell lines (**Figure 4b**). For the EGFR inhibitors erlotinib and sapitinib, we observed that MES models were slightly more sensitive compared to ADRN-like models (**Supplemental Figure 8a**), whereas PDGFR inhibition did not show subtype-specific cytotoxicity (**Supplemental Figure 8b**).

**Figure 4.**
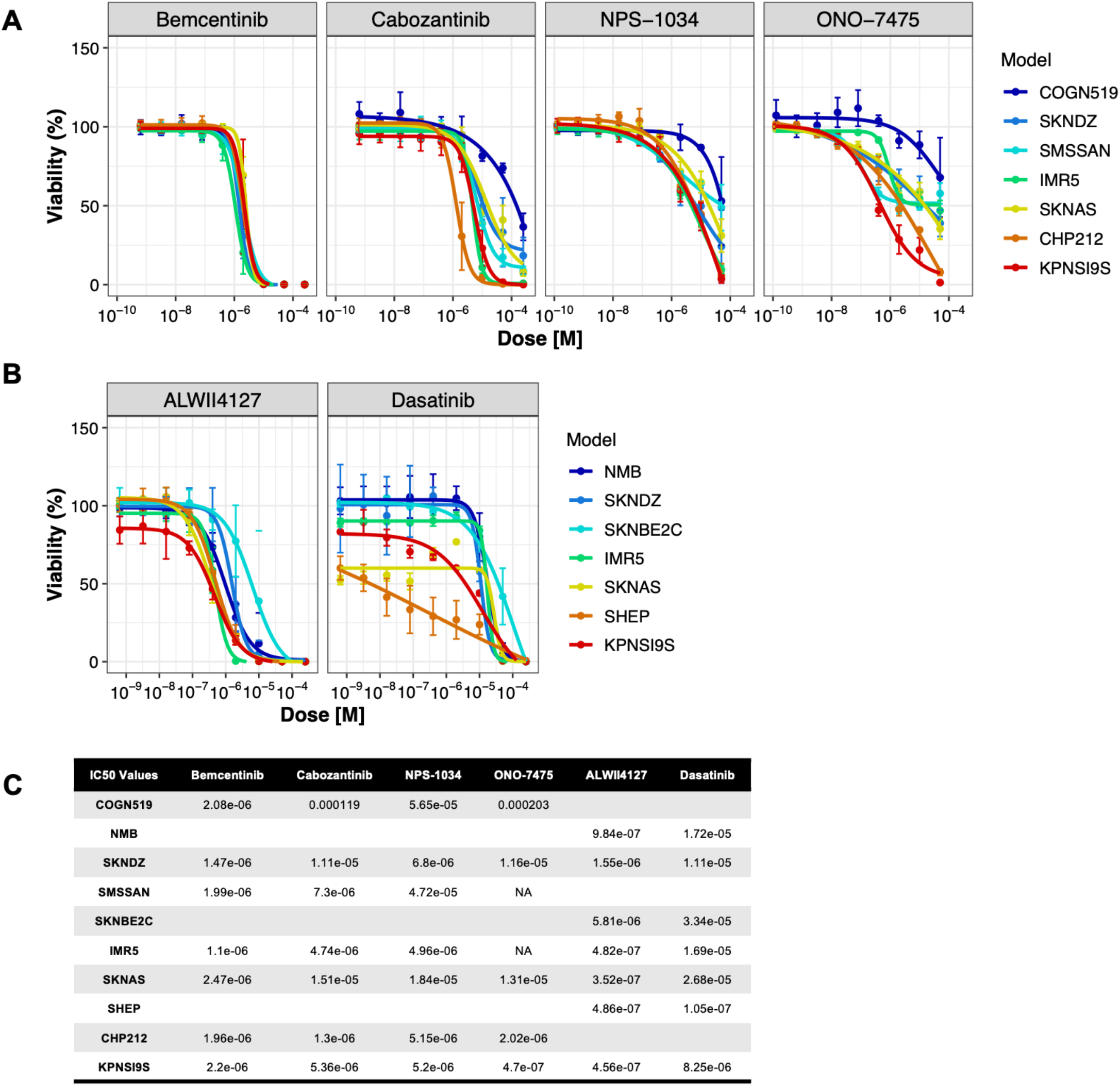
RTK-targeted small molecule inhibitors show specificity towards MES neuroblastoma cell line models. A-C) Cytotoxicity of small molecule inhibitors targeting (A) AXL and (B) EPHA2 / multiple tyrosine kinases across a panel of ADRN and MES neuroblastoma cell lines with (C) associated IC50 values (M). Viability was measured with CellTiter-Glo2.0 after 3 days of exposure to each inhibitor in 2-3 independent experiments.

*AXL-specific antibody-drug conjugate (ADCT-601) shows anti-tumor activity.* Because we posit that AXL has both tumor cell intrinsic and extrinsic functions in mediating a MES cell state, and it appears that AXL signaling alone is not a major oncogenic drive of the MES cell state, we explored ADCT-601, an AXL-targeted ADC, which contains a potent pyrrolobenzodiazepine dimer toxin to explore MES-specific therapeutic targeting. We evaluated the activity of ADCT-601 and isotype control ADC B12-PL1601 (targeting HIV envelop protein gp120) across a panel of neuroblastoma cell lines. In an ADRN model (IMR5) and a MES-derived model with *AXL* genetically deleted (CHP212-sgAXL), we observed no difference between the ADCT-601 and B12-PL1601 (**Figure 5a**). Conversely, MES-like models with cell-surface AXL (**Figures 3d** **and Supplemental Figure 6d-e**) were more sensitive to ADCT-601 compared to B12-PL1601 (**Figure 5a**). We calculated ADCT-601 IC50 values of 6.52pM in KPNSI9s and 74.5nM in CRISPR control model CHP212-sgCHR2 (**Figure 5b**). The MES model SHEP was generally less sensitive to these agents (IC50s: ADCT-601 at 1.35nM and B12-PL1601 at 11.7nM). Our observed IC50 values for ADCT-601 are within previously published ranges (20pM – 2.2nM).^50^

**Figure 5.**
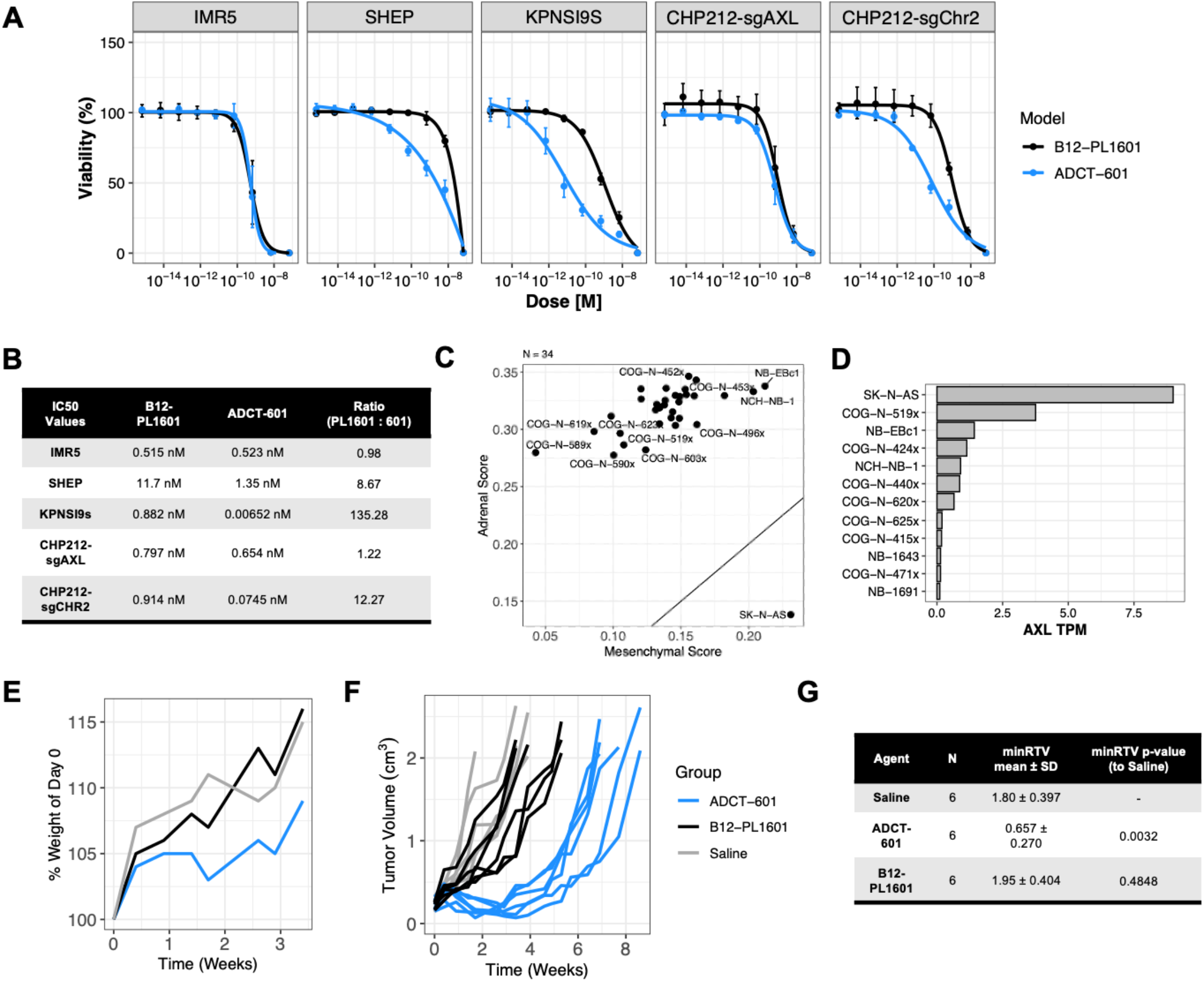
AXL-specific antibody-drug conjugate (ADCT-601) demonstrates activity *in vitro* and *in vivo*. A) Cellular viability and (B) IC50 values after 5 days of treatment with Control ADC (B12-PL1601) or AXL-targeted ADC (ADCT-601). Data was collected from three independent experiments. Ratio reported is B12- PL1601:ADCT-601. C) ADR/MES signature scores and (D) AXL mRNA expression in patient-derived and cell-line-derived xenograft models. E) Mouse weights after treatment (day 0) as a percent of day 0. F) Tumor volumes after treatment (day 0) of individual mice. Treatments are Saline, Control ADC (B12-PL1601, 1mg/kg dose), or AXL-targeted ADC (ADCT-601, 1mg/kg dose). N=6 mice per treatment arm. G) Minimum relative tumor volume (minRTV) mean and standard deviation (SD) per treatment group with associated statistics using pairwise Wilcoxon Rank-Sum.

We next extended our *in vitro* studies to patient-derived and cell-line-derived xenograft models. Only one patient- or cell line-derived xenograft (CDX), SKNAS, was MES-dominant and expressed *AXL* (**Figure 5c-d**). *In vivo* studies with SKNAS (N=6/arm) and the ADRN COGN519x PDX (N=3/arm) show that a single dose of ADCT- 601 at 1mg/kg was well-tolerated (**Figure 5e, Supplemental Figure 9a**). In SKNAS-bearing animals, we observed a significant decrease in minimum relative tumor volume (minRTV) compared to the saline control (p = 0.0032; **Figure 5f-g**). Anti-tumor activity with ADCT-601 was not seen in the ADRN COGN519x model (**Supplemental Figure 9b**). Altogether, our data supports the further development of AXL-targeted ADCs in MES-dominant neuroblastoma models.

## Discussion

Neuroblastoma is a heterogeneous disease in terms of clinical manifestations, natural history, and response to therapy. While somatic genomic heterogeneity explains in part clinical diversity and therapy resistance,^59,60^ it has been known for almost 50 years that neuroblastoma cells in culture can assume vastly different cell states.^26^ More recently, the epigenetic basis of this observation has been defined,^28,29,61^ but the relevance to patients and therapy remains largely theoretical. Here, we characterized patient tumors by their dominant signature (ADRN or MES) to evaluate the expression of proven and candidate therapeutic targets in the admixed cells that reside in neuroblastomas. We extended prior observations that the MES subtype is more immunogenic by observing an association with the inflammatory response^52,53^. These genes also show subtype-specific histone marks at these loci and forced transdifferentiation from ADRN to MES is associated with significantly increased expression. This suggests that MES cells may be vulnerable to immunotherapeutic approaches, perhaps targeting cancer-specific peptides presented on common HLA allotypes,^62^ or cell surface proteins enriched in this state. It is notable, however, that HLA-E, which inhibits NK and CD8^+^ cytotoxic T cells^63,64^, expression is also increased in MES cells and may confer immune evasion. Whether blocking HLA-E:NKG2A interaction with specific antibodies^65,66^ renders the MES cells more susceptible to immunotherapy remains to be established.

Our study underscores the importance of neuroblastoma subtype heterogeneity and its implications on targeted therapies. We show that multiple immunotherapeutic targets in clinical development are ADRN- predominant and are expressed at significantly lower levels, or even absent, in MES-like tumors and cell lines. These findings are supported by ADRN-to-MES transdifferentiation experiments in cell line models and evaluation of the epigenome in parental cell lines. In contrast, *CD276* and *L1CAM* show relatively stable expression in ADRN and MES cells. In our recent preclinical study using a CD276-targeted ADC (m276-SL- PBD), we observed an objective response rate of 93% in the neuroblastoma cell line-derived and patient-derived xenografts.^19^ Importantly, all neuroblastoma PDXs assessed display a strong ADRN signature, so the efficacy of CD276-targeted agents, as well as other preclinical molecules, remains to be determined in MES-dominant neuroblastomas. While antigen loss is a known mechanisms of escape from many immunotherapies, our data suggest that this might largely occur by epigenetic mediated cell state transitions in neuroblastoma and this needs to be considered in ongoing and future clinical trials.

We identified several candidate MES-specific receptor tyrosine kinases, including *AXL, EPHA2, EGFR*, and *PDGFRA/B*. Several of these genes have been described as important mediators of cancer cell resistance to targeted therapies.^56–58,67^ Recently, Noronha et al. describe increased signaling through the GAS6-AXL signaling axis in drug-tolerate persister cells, which can mediate resistance to RTK inhibitors in *EGFR*-mutant lung cancer models.^68^ We showed enrichment of these proteins in MES cell lines and observed upregulation of these genes after ADRN-to-MES transdifferentiation. However, despite AXL being a differently expressed gene in MES- dominant tumor samples and cell line models, here we show that it is neither necessary for MES subtype maintenance nor sufficient to transdifferentiate ADRN cells to a MES-like phenotype. Thus, the potential role of AXL RTK inhibitors to target MES cell state in this disease requires further exploration, but it is of interest that the multi-RTK inhibitor dasatinib has been used in a metronomic regimen for disease palliation.^69^

Finally, we assessed the activity of ADCT-601, a pyrrolobenzodiazepine dimer-based AXL-targeted ADC, in neuroblastoma cell lines and CDX/PDX models. We show that a single dose of ADCT-601 is well-tolerated and demonstrates significant anti-tumor activity in a MES CDX model. AXL-targeted antibody-drug conjugates have been evaluated in non-small cell lung cancer,^70^ soft tissue sarcoma,^71^ and melanoma,^72^ in combination with small molecule inhibitors or immune checkpoint blockade,^73^ and Phase I clinical trials with ADCT-601 are ongoing (NCT05389462). As a pan-cancer mesenchymal marker, AXL upregulation can be observed after treatment with targeted therapy.^58^ Thus, combination strategies with AXL-targeted ADCs aim to prevent resistance by targeting therapy-induced heterogeneity.^72^ In experimental neuroblastoma models, AXL has been reported to contribute to therapy resistance to ALK inhibitors.^57^ Future studies evaluating various treatment schedules of ADCT-601 in additional MES-dominant models will be essential to further determine the efficacy of this compound. Additionally, studies incorporating heterogenous tumors with combination strategies will be necessary to credential AXL as an effective therapy for the elimination of MES-specific neuroblastoma tumor cells and/or the treatment-resistant neuroblastomas.

As most high-risk neuroblastomas are ADRN-dominant, successful clinical developments have focused on targeting molecules abundant in this subtype. However, studies investigating neuroblastoma heterogeneity show the plasticity of neuroblastoma cells and consequently, the ADRN-specific molecules of clinical interest. Therefore, it is critical to further profile this subset of neuroblastoma cells and develop approaches to targets the cells able to persistent ADRN-targeted modalities. As more targeted therapies are brought forward to the clinic, serial monitoring of tumor specimens will be essential to understand the temporal heterogeneity in relation to treatment response and allow clinicians to identify and treat emerging resistance.

## Acknowledgements

This work was supported by a St. Baldrick’s-Stand Up To Cancer Pediatric Dream Team Translational Research Grant (SU2C-AACR-DT2727). Stand Up to Cancer is a program of the Entertainment Industry Foundation administered by the American Association for Cancer Research (JMM). This work was also supported by the following NIH grants: P01 CA217959 (JMM, YPM and KS), U54 CA232568 as part of the Beau Biden Cancer Moonshot Program (JMM), R35 CA220500 (JMM), R35 CA210030 (KS), F31 CA239424 (NMK), F32 CA261035

(NWM), and K08 CA266914 (AJW). This work was also supported by a Department of Defense Career Development Award W81XWH2210474 (AJW) and the Giulio D’Angio Endowed Chair (JMM).

## Authorship

Study Conception: NMK, JMM Experimental Design & Execution: NMK, MO, NWM, MT, DG Data Curation: NMK, NWM Computational Support: AF, LG Experimental Support: AW, FZ, PB, CD, YM, KS Manuscript Draft: NMK Manuscript Review: all authors

## Data Availability

Code and data produced will be made available upon reasonable request.

---

**S. Figure 1.**
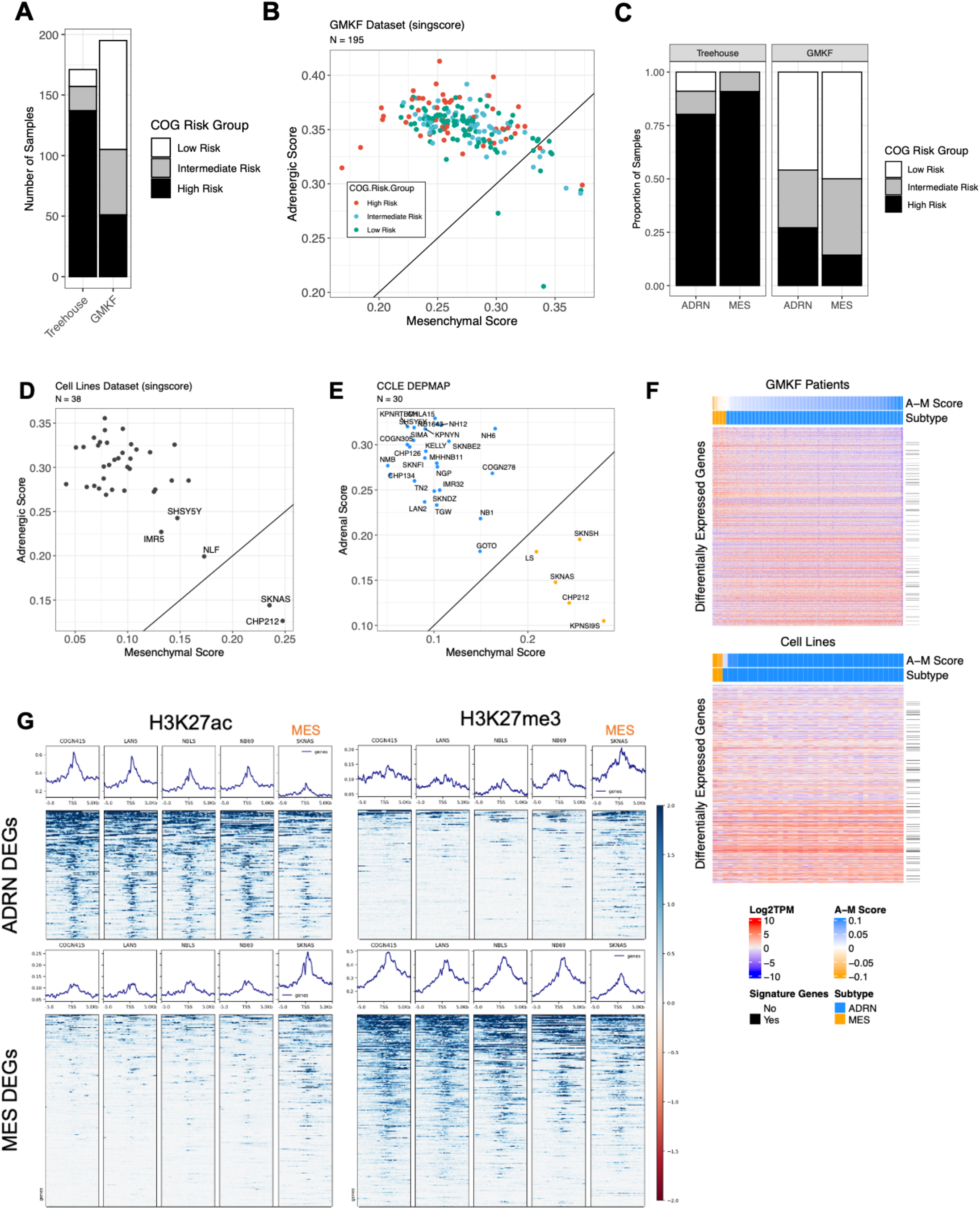
ADRN- and MES-dominant patient samples and cell lines are characterized by differentially expressed genes driven by subtype-specific enhancers. S. Figure 1 Legend: A) Distribution of COG Risk Group across Treehouse and GMKF datasets, (B) subset by a ADRN and MES. C-E) ADRN and MES scores calculated by single-sample gene set enrichment analysis (singscore) in (C) GMKF patient samples and (D-E) neuroblastoma cell line models. E-F) Heatmaps of top 500 MES and ADRN differentially expressed genes (from Treehouse) in (E) GMKF patient samples and (F) neuroblastoma cell line models. G) Histone ChIP-sequencing (H3K27ac or H3K27me3) in ADRN or MES neuroblastoma cell lines at ADRN and MES differentially expressed genes.

**Supplemental Figure 2.**
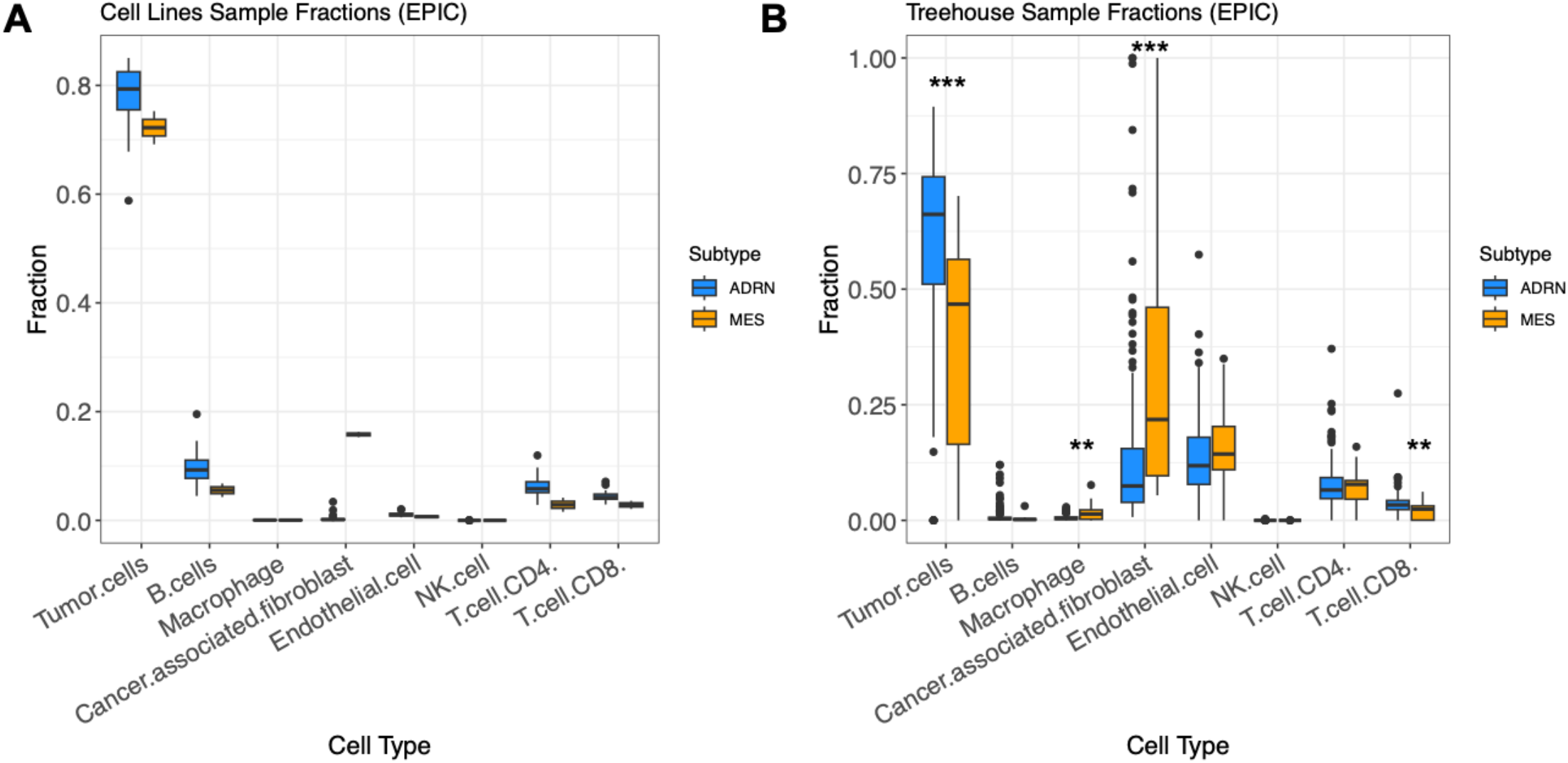
Estimated tumor and immune cell fractions in bulk RNA-sequencing datasets. S. Figure 2 Legend: A-B) Tumor and immune cell fractions predicted with deconvolution algorithm EPIC in (A) neuroblastoma cell lines and (B) Treehouse patient bulk RNA-sequencing datasets. Statistics are as follows: * p<0.5, ** p<0.01, *** p<0.001.

**Supplemental Figure 3.**
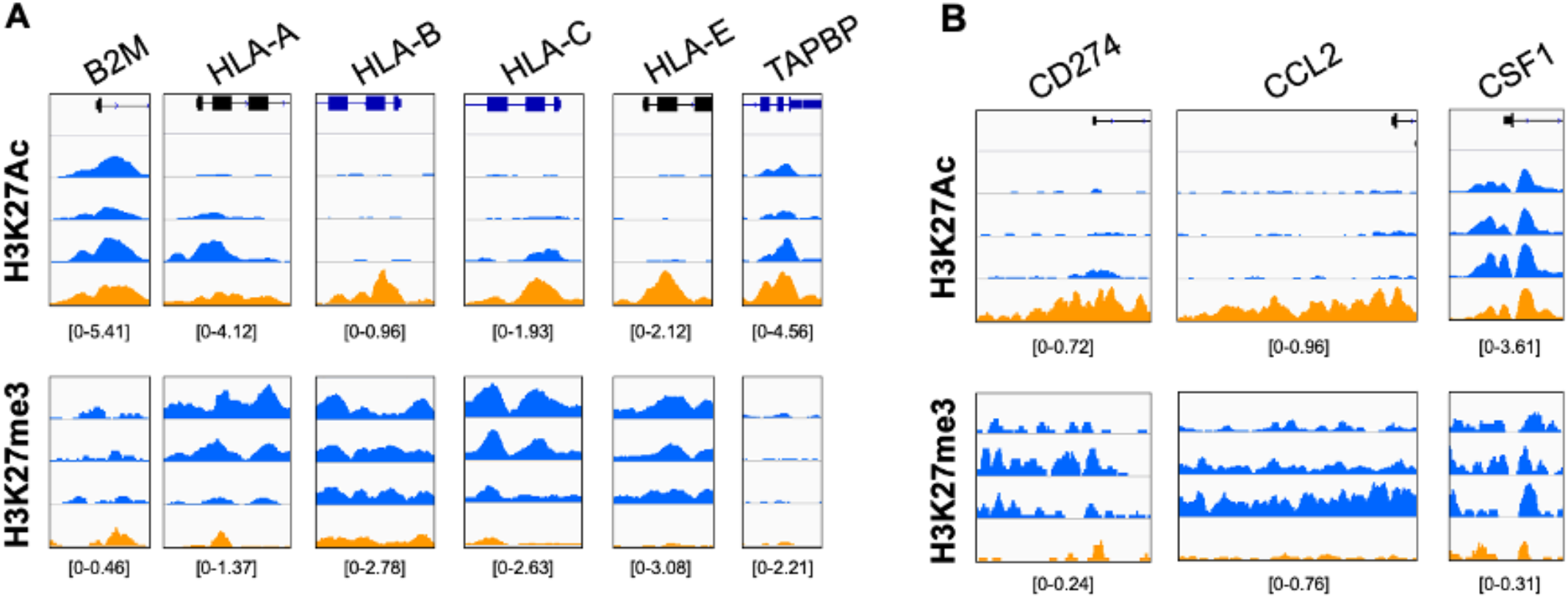
Epigenetic regulation of MHC processing/presentation genes and immunomodulatory genes. S. Figure 3 Legend: A-B) Histone ChIP-sequencing peaks at target loci representing H3K27ac (enhancer, top) and H3K27me3 (repressive, bottom) marks in COGN415, KELLY, SKNBE2C, and SKNAS models. Colors represent ADRN (blue) and MES (orange) cell lines.

**Supplemental Figure 4.**
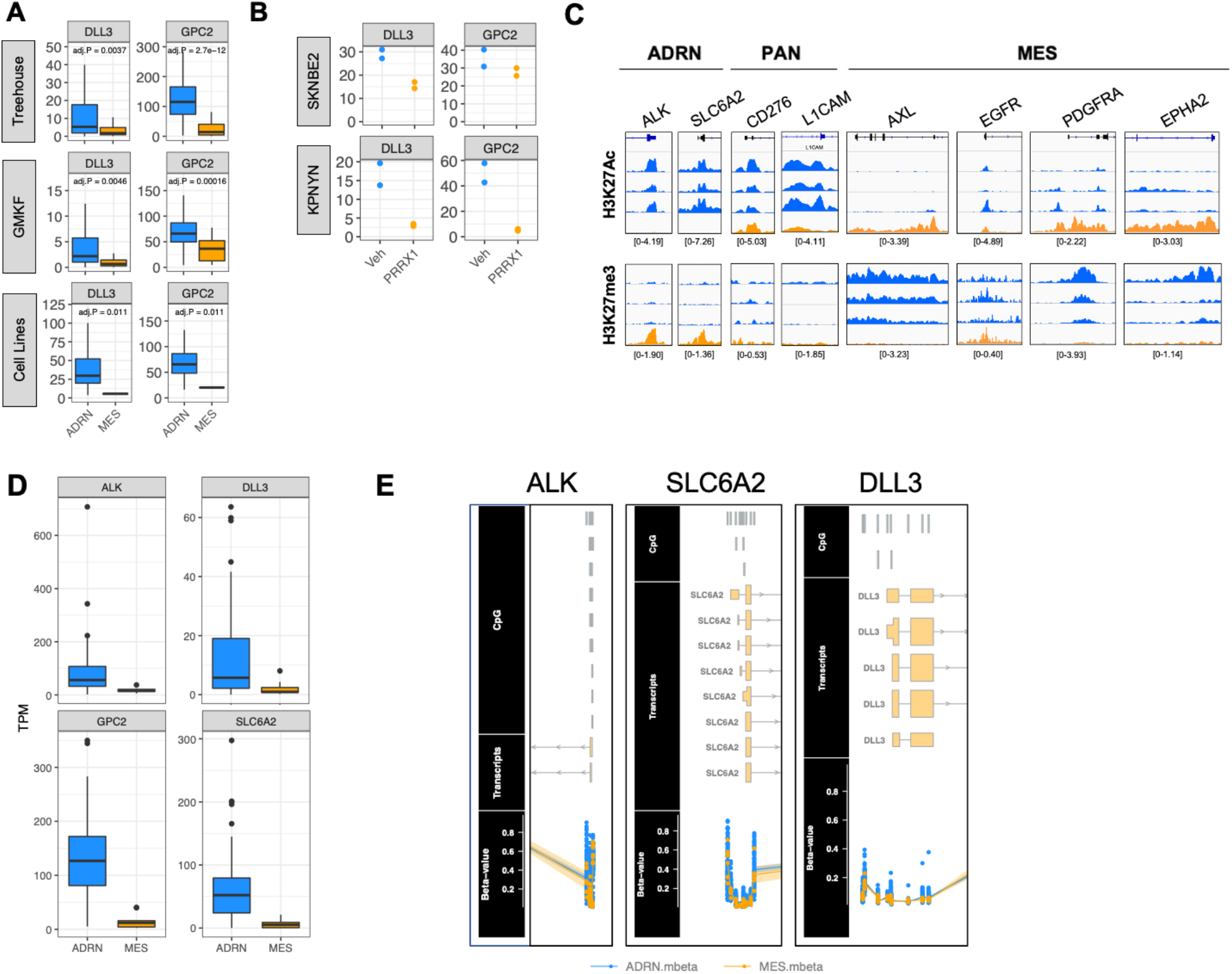
Differential expression and epigenetic regulation of candidate subtype-specific and pan-subtype targets. S. Figure 4 Legend: A-B) RNA expression of preclinical ADRN-specific targets, DLL3 and GPC2, in (A) patient datasets, cell lines, and (B) ADRN-to-MES transdifferentiation models. B) Histone ChIP- sequencing peaks at target loci representing H3K27ac (enhancer, top) and H3K27me3 (repressive, bottom) marks in COGN415, KELLY, SKNBE2C, and SKNAS models. D) RNA expression of ADRN-specific targets in subset of patient samples with methylation sequencing data. E). mBeta scores in ADRN-dominant and MES-dominant patients at the ALK, SLC6A2, and DLL3 loci.

**Supplemental Figure 5.**
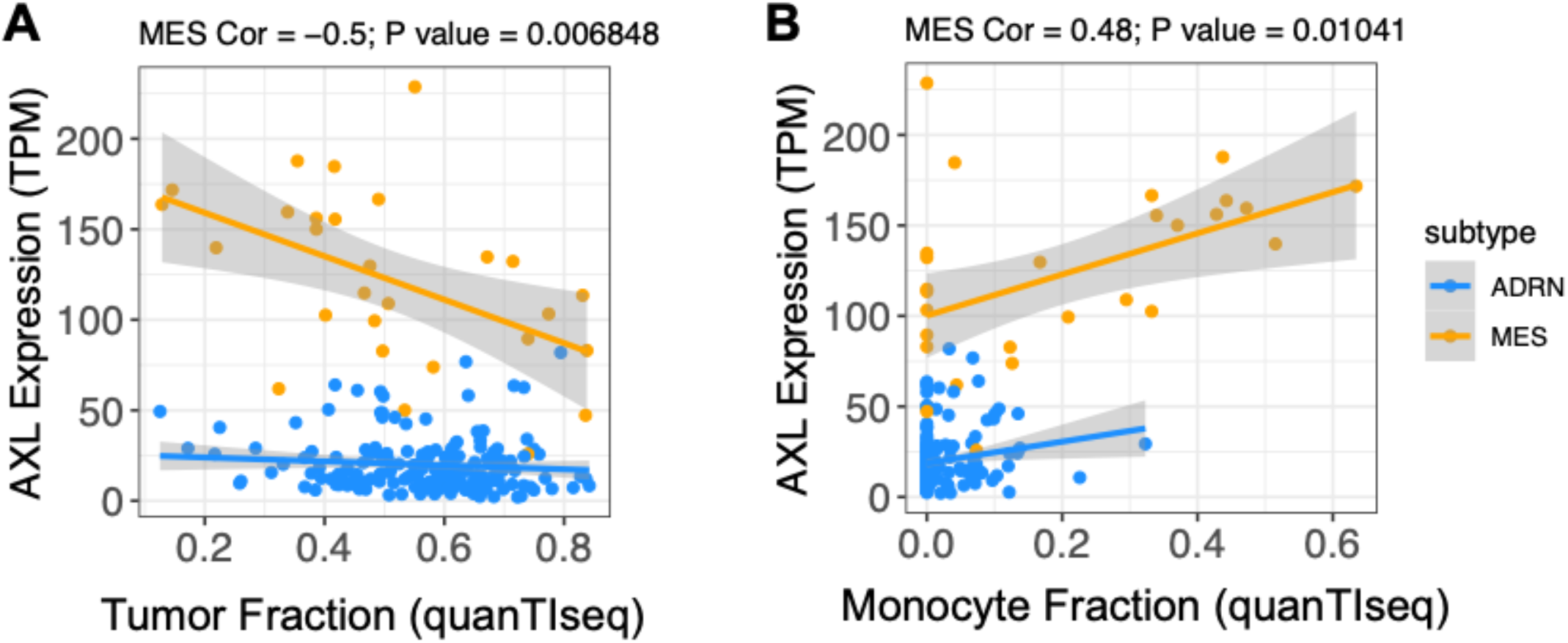
Correlation between predicted fraction (quanTIseq) and AXL mRNA expression. S. Figure 5 Legend: A-B) AXL mRNA expression (TPM) vs. (A) tumor and (B) monocyte fractions predicted by quanTIseq. Spearman correlations presented are only for MES samples.

**Supplemental Figure 6.**
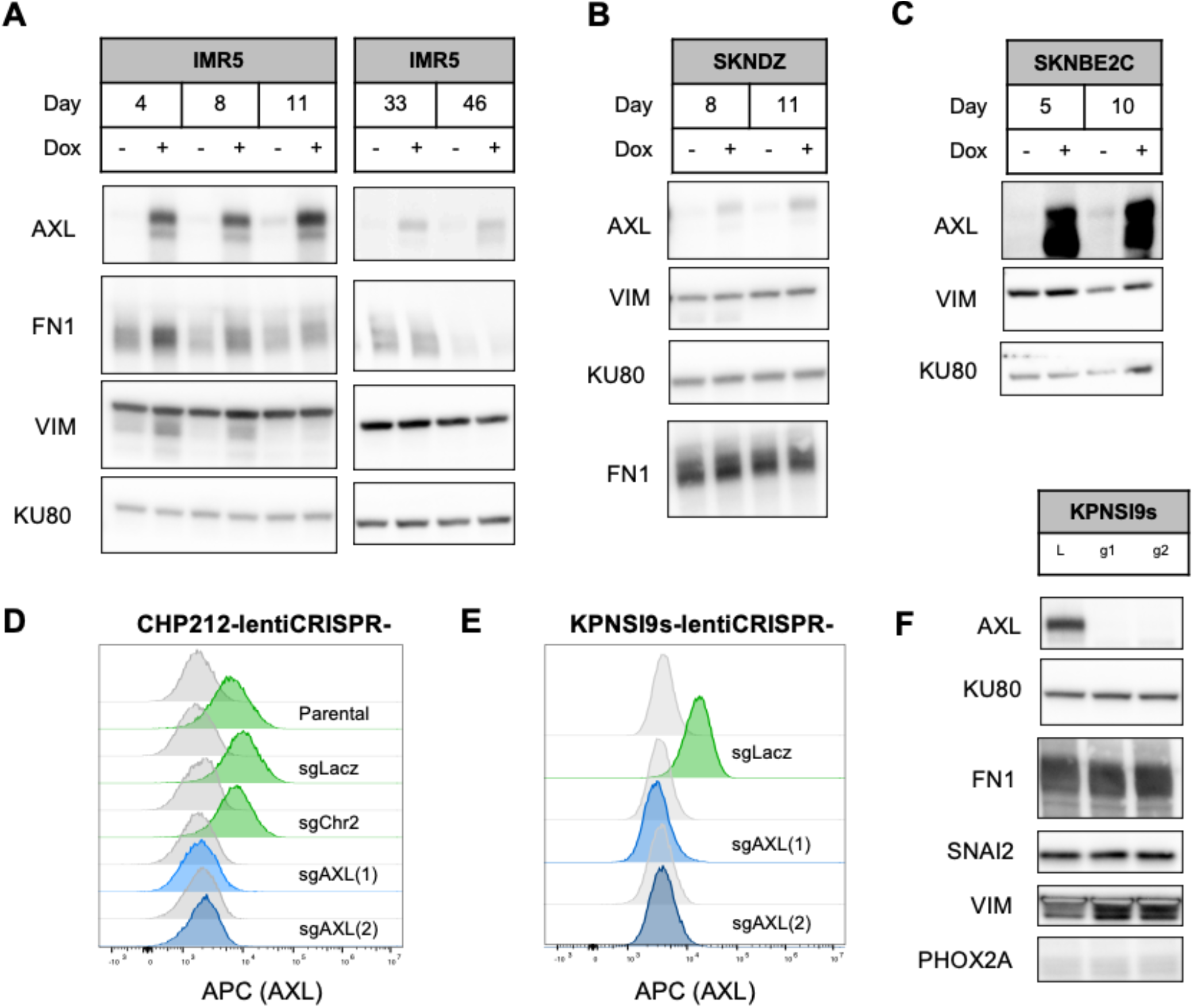
AXL is not sufficient to change, nor necessary to maintain the MES phenotype. S. Figure 6 Legend: A-C) AXL overexpression was achieved by subsequent transduction of ADRN parental cell lines (IMR5, SKNDZ, SKNBE2C) with lentiviral particles containing the tet repressor protein and then the AXL transgene driven by a CMV/TO promoter. Cells were cultured in the presence of DMSO or doxycycline (1ug/mL) for up to 46 days and assayed for both MES and ADRN markers. D-E) AXL expression measured by flow cytometry after knockout of AXL in the MES models (D) CHP212 and (E) KPNSI9s with the pLentiCRISPRv2 system using two sgRNAs and a control sgRNA targeting LACZ and/or Chr2.2. F) Expression of MES and ADRN markers in KPNSI9s models after AXL knockout.

**Supplemental Figure 7.**
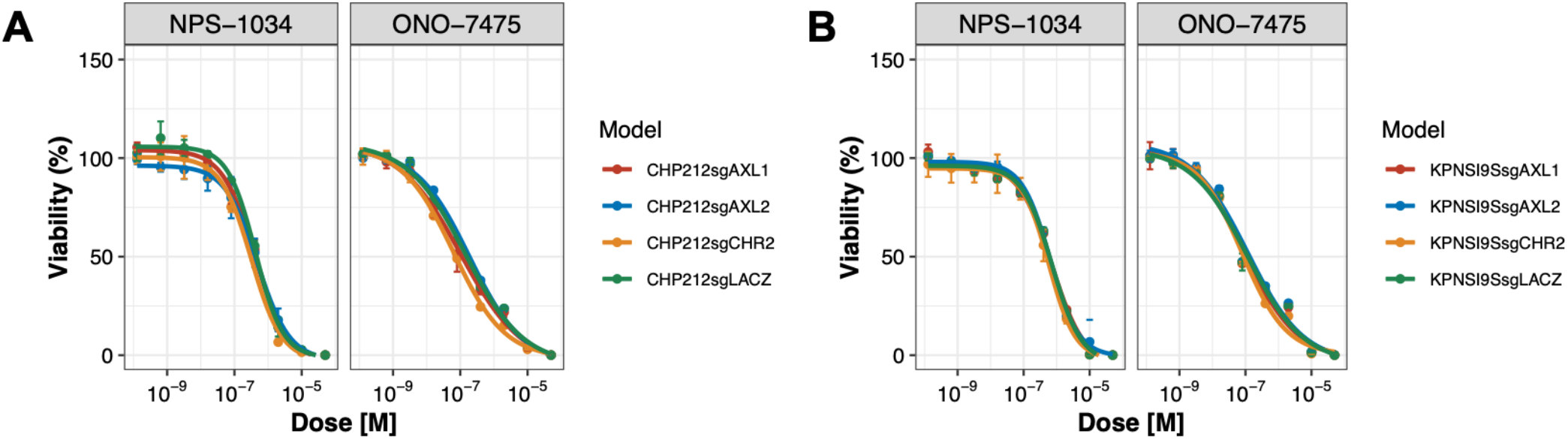
AXL inhibitors do not exclusively target AXL. S. Figure 7 Legend: A-B) Cytotoxicity of small molecule AXL inhibitors (NPS-1034 and ONO-7475) in (A) CHP212 and (B) KPNSI9s CRISPR knockout controls (sgLacz, sgCHR2) or AXL knockout (sgAXL1, sgAXL2) models. Viability was measured with CellTiter-Glo2.0 after 3 days of exposure to each inhibitor. Replicates are technical.

**Supplemental Figure 8.**
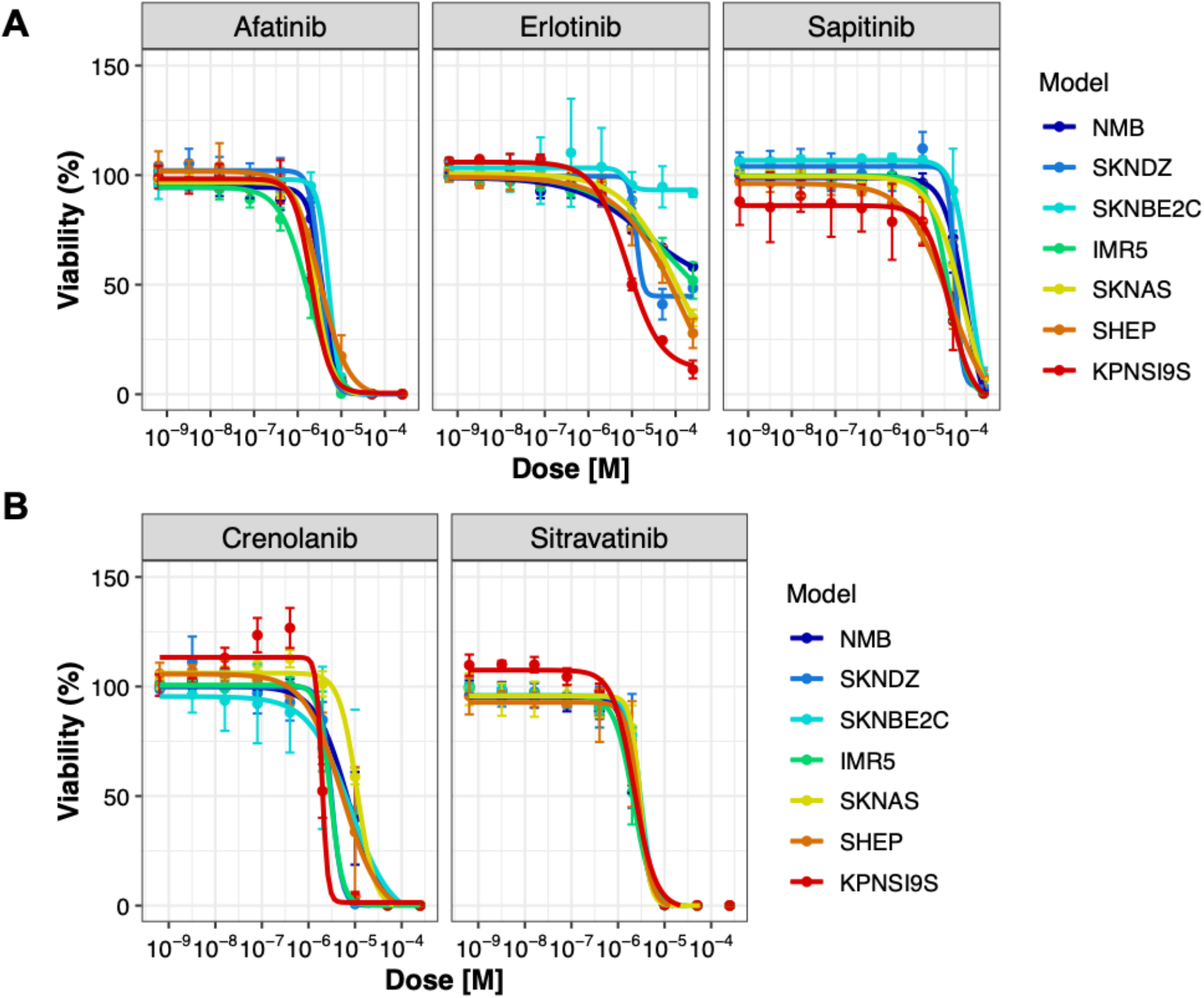
EGFR- and PDGFR-targeted small molecule inhibitors show modest specificity towards MES neuroblastoma cell line models.| S. Figure 8 Legend: A-B) Cytotoxicity of small molecule inhibitors targeting (A) EGFR and (B) PDGFRA/B across a panel of ADRN and MES neuroblastoma cell lines. Viability was measured with CellTiter-Glo2.0 after 3 days of exposure to each inhibitor after 2-3 independent experiments.

**Supplemental Figure 9.**
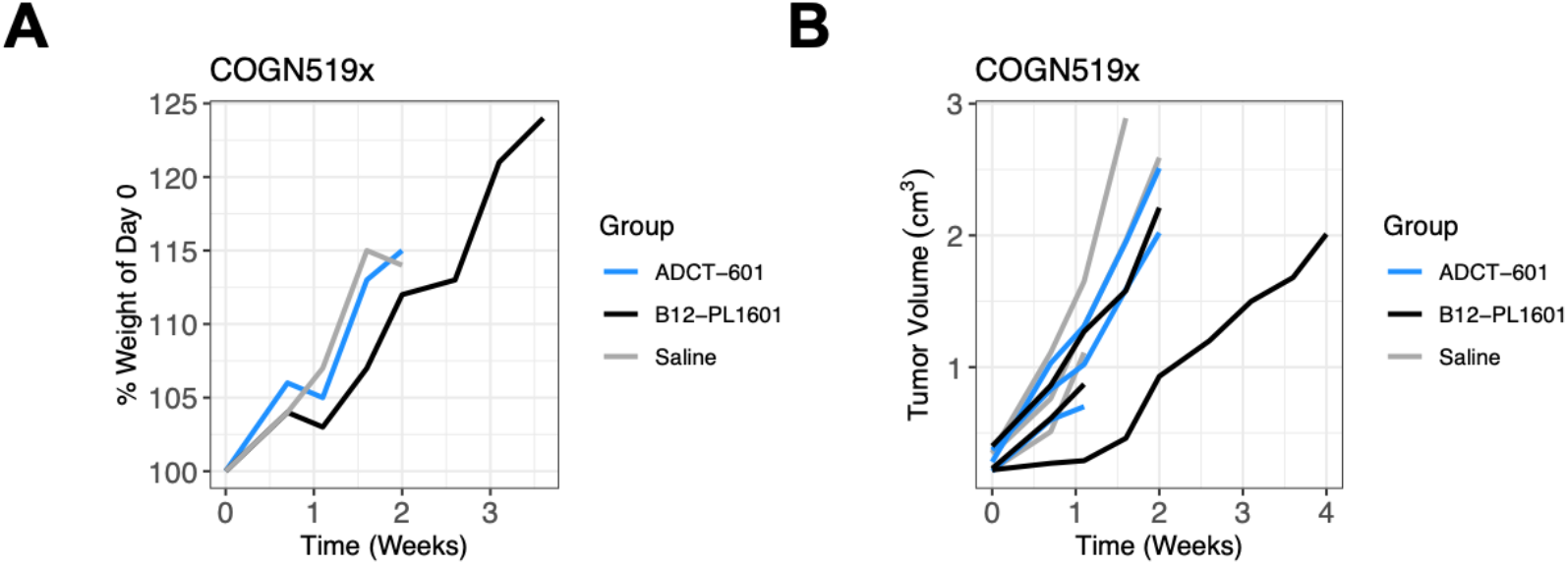
ADCT-601 is only active in MES-dominant model, SKNAS. S. Figure 9 Legend: A) Mouse weights after treatment (day 0) as a percent of day 0. B) Tumor volumes after treatment (day 0) of individual mice. Treatments are Saline, Control ADC (B12- PL1601, 1mg/kg dose), or AXL-targeted ADC (ADCT-601, 1mg/kg dose). N=2-3 animals per treatment arm.

## References

1. Maris JM. Recent advances in neuroblastoma. N Engl J Med. 2010; 362(23):2202–2211.

2. Matthay KK, Maris JM, Schleiermacher G, et al. Neuroblastoma. Nat Rev Dis Primers. 2016; 2:16078.

3. Yu AL, Gilman AL, Ozkaynak MF, et al. Anti-GD2 antibody with GM-CSF, interleukin-2, and isotretinoin for neuroblastoma. N Engl J Med. 2010; 363(14):1324–1334.

4. Yu AL, Gilman AL, Ozkaynak MF, et al. Long-Term Follow-up of a Phase III Study of ch14.18 (Dinutuximab) + Cytokine Immunotherapy in Children with High-Risk Neuroblastoma: COG Study ANBL0032. Clin Cancer Res. 2021.

5. Ladenstein R, Potschger U, Valteau-Couanet D, et al. Interleukin 2 with anti-GD2 antibody ch14.18/CHO (dinutuximab beta) in patients with high-risk neuroblastoma (HR-NBL1/SIOPEN): a multicentre, randomised, phase 3 trial. Lancet Oncol. 2018; 19(12):1617–1629.

6. Kushner BH, Cheung IY, Modak S, Basu EM, Roberts SS, Cheung NK. Humanized 3F8 Anti-GD2 Monoclonal Antibody Dosing With Granulocyte-Macrophage Colony-Stimulating Factor in Patients With Resistant Neuroblastoma: A Phase 1 Clinical Trial. JAMA Oncol. 2018; 4(12):1729–1735.

7. Mabe NW, Huang M, Dalton GN, et al. Transition to a mesenchymal state in neuroblastoma confers resistance to anti-GD2 antibody via reduced expression of ST8SIA1. Nat Cancer. 2022.

8. Del Bufalo F, De Angelis B, Caruana I, et al. GD2-CART01 for Relapsed or Refractory High-Risk Neuroblastoma. N Engl J Med. 2023; 388(14):1284–1295.

9. Majzner RG, Ramakrishna S, Yeom KW, et al. GD2-CAR T cell therapy for H3K27M-mutated diffuse midline gliomas. Nature. 2022.

10. Infarinato NR, Park JH, Krytska K, et al. The ALK/ROS1 inhibitor PF-06463922 overcomes primary resistance to crizotinib in ALK-driven neuroblastoma. Cancer Discov. 2015.

11. Berko ER, Witek GM, Matkar S, et al. Circulating tumor DNA reveals mechanisms of lorlatinib resistance in patients with relapsed/refractory ALK-driven neuroblastoma. Nat Commun. 2023; 14(1):2601.

12. Goldsmith KC, Park JR, Kayser K, et al. Lorlatinib with or without chemotherapy in ALK-driven refractory/relapsed neuroblastoma: phase 1 trial results. Nat Med. 2023.

13. Walker AJ, Majzner RG, Zhang L, et al. Tumor Antigen and Receptor Densities Regulate Efficacy of a Chimeric Antigen Receptor Targeting Anaplastic Lymphoma Kinase. Mol Ther. 2017; 25(9):2189–2201.

14. Bergaggio E, Tai WT, Aroldi A, et al. ALK inhibitors increase ALK expression and sensitize neuroblastoma cells to ALK.CAR-T cells. Cancer Cell. 2023; 41(12):2100–2116 e2110.

15. Sano R, Krytska K, Larmour CE, et al. An antibody-drug conjugate directed to the ALK receptor demonstrates efficacy in preclinical models of neuroblastoma. Sci Transl Med. 2019; 11(483).

16. Hong H, Stastny M, Brown C, et al. Diverse solid tumors expressing a restricted epitope of L1-CAM can be targeted by chimeric antigen receptor redirected T lymphocytes. J Immunother. 2014; 37(2):93–104.

17. Kunkele A, Taraseviciute A, Finn LS, et al. Preclinical Assessment of CD171-Directed CAR T-cell Adoptive Therapy for Childhood Neuroblastoma: CE7 Epitope Target Safety and Product Manufacturing Feasibility. Clin Cancer Res. 2017; 23(2):466–477.

18. Moghimi B, Muthugounder S, Jambon S, et al. Preclinical assessment of the efficacy and specificity of GD2-B7H3 SynNotch CAR-T in metastatic neuroblastoma. Nat Commun. 2021; 12(1):511.

19. Kendsersky NM, Lindsay JM, Kolb EA, et al. The B7-H3-targeting antibody-drug conjugate m276-SL- PBD is potently effective against pediatric cancer preclinical solid tumor models. Clin Cancer Res. 2021.

20. Theruvath J, Sotillo E, Mount CW, et al. Locoregionally administered B7-H3-targeted CAR T cells for treatment of atypical teratoid/rhabdoid tumors. Nat Med. 2020; 26(5):712–719.

21. Vitanza NA, Wilson AL, Huang W, et al. Intraventricular B7-H3 CAR T Cells for Diffuse Intrinsic Pontine Glioma: Preliminary First-in-Human Bioactivity and Safety. Cancer Discov. 2023; 13(1):114–131.

22. Bosse KR, Raman P, Zhu Z, et al. Identification of GPC2 as an Oncoprotein and Candidate Immunotherapeutic Target in High-Risk Neuroblastoma. Cancer Cell. 2017; 32(3):295–309 e212.

23. Heitzeneder S, Bosse KR, Zhu Z, et al. GPC2-CAR T cells tuned for low antigen density mediate potent activity against neuroblastoma without toxicity. Cancer Cell. 2022; 40(1):53–69 e59.

24. Raman S, Buongervino SN, Lane MV, et al. A GPC2 antibody-drug conjugate is efficacious against neuroblastoma and small-cell lung cancer via binding a conformational epitope. Cell Rep Med. 2021; 2(7):100344.

25. Li N, Torres MB, Spetz MR, et al. CAR T cells targeting tumor-associated exons of glypican 2 regress neuroblastoma in mice. Cell Rep Med. 2021; 2(6):100297.

26. Biedler JL, Helson L, Spengler BA. Morphology and growth, tumorigenicity, and cytogenetics of human neuroblastoma cells in continuous culture. Cancer Research. 1973; 33(11):2643–2652.

27. Ross RA, Biedler JL, Spengler BA. A role for distinct cell types in determining malignancy in human neuroblastoma cell lines and tumors. Cancer Lett. 2003; 197(1-2):35–39.

28. Boeva V, Louis-Brennetot C, Peltier A, et al. Heterogeneity of neuroblastoma cell identity defined by transcriptional circuitries. Nat Genet. 2017; 49(9):1408–1413.

29. van Groningen T, Koster J, Valentijn LJ, et al. Neuroblastoma is composed of two super-enhancer-associated differentiation states. Nat Genet. 2017; 49(8):1261–1266.

30. van Groningen T, Akogul N, Westerhout EM, et al. A NOTCH feed-forward loop drives reprogramming from adrenergic to mesenchymal state in neuroblastoma. Nat Commun. 2019; 10(1):1530.

31. Wang L, Tan TK, Durbin AD, et al. ASCL1 is a MYCN- and LMO1-dependent member of the adrenergic neuroblastoma core regulatory circuitry. Nat Commun. 2019; 10(1):5622.

32. Durbin AD, Zimmerman MW, Dharia NV, et al. Selective gene dependencies in MYCN-amplified neuroblastoma include the core transcriptional regulatory circuitry. Nat Genet. 2018; 50(9):1240–1246.

33. Decaesteker B, Denecker G, Van Neste C, et al. TBX2 is a neuroblastoma core regulatory circuitry component enhancing MYCN/FOXM1 reactivation of DREAM targets. Nat Commun. 2018; 9(1):4866.

34. Soldatov R, Kaucka M, Kastriti ME, et al. Spatiotemporal structure of cell fate decisions in murine neural crest. Science. 2019; 364(6444).

35. Robinson MD, McCarthy DJ, Smyth GK. edgeR: a Bioconductor package for differential expression analysis of digital gene expression data. Bioinformatics. 2010; 26(1):139–140.

36. McCarthy DJ, Chen Y, Smyth GK. Differential expression analysis of multifactor RNA-Seq experiments with respect to biological variation. Nucleic Acids Res. 2012; 40(10):4288–4297.

37. Harenza JL, Diamond MA, Adams RN, et al. Transcriptomic profiling of 39 commonly-used neuroblastoma cell lines. Sci Data. 2017; 4:170033.

38. Rokita JL, Rathi KS, Cardenas MF, et al. Genomic Profiling of Childhood Tumor Patient-Derived Xenograft Models to Enable Rational Clinical Trial Design. Cell reports. 2019; 29(6):1675–1689 e1679.

39. Foroutan M, Bhuva DD, Lyu R, Horan K, Cursons J, Davis MJ. Single sample scoring of molecular phenotypes. BMC Bioinformatics. 2018; 19(1):404.

40. Law CW, Chen Y, Shi W, Smyth GK. voom: Precision weights unlock linear model analysis tools for RNA-seq read counts. Genome Biol. 2014; 15(2):R29.

41. Ritchie ME, Phipson B, Wu D, et al. limma powers differential expression analyses for RNA-sequencing and microarray studies. Nucleic Acids Res. 2015; 43(7):e47.

42. Barry E, Walsh JA, Weinrich SL, et al. Navigating the Regulatory Landscape to Develop Pediatric Oncology Drugs: Expert Opinion Recommendations. Paediatr Drugs. 2021; 23(4):381–394.

43. Finotello F, Mayer C, Plattner C, et al. Molecular and pharmacological modulators of the tumor immune contexture revealed by deconvolution of RNA-seq data. Genome Med. 2019; 11(1):34.

44. Racle J, Gfeller D. EPIC: A Tool to Estimate the Proportions of Different Cell Types from Bulk Gene Expression Data. Methods Mol Biol. 2020; 2120:233–248.

45. Upton K, Modi A, Patel K, et al. Epigenomic profiling of neuroblastoma cell lines. Sci Data. 2020; 7(1):116.

46. Lalchungnunga H, Hao W, Maris JM, et al. Genome wide DNA methylation analysis identifies novel molecular subgroups and predicts survival in neuroblastoma. Br J Cancer. 2022.

47. Subramanian A, Tamayo P, Mootha VK, et al. Gene set enrichment analysis: a knowledge-based approach for interpreting genome-wide expression profiles. Proc Natl Acad Sci U S A. 2005; 102(43):15545–15550.

48. Liberzon A, Birger C, Thorvaldsdottir H, Ghandi M, Mesirov JP, Tamayo P. The Molecular Signatures Database (MSigDB) hallmark gene set collection. Cell Syst. 2015; 1(6):417–425.

49. Sanjana NE, Shalem O, Zhang F. Improved vectors and genome-wide libraries for CRISPR screening. Nat Methods. 2014; 11(8):783–784.

50. Zammarchi F, Havenith KE, Chivers S, et al. Preclinical Development of ADCT-601, a Novel Pyrrolobenzodiazepine Dimer-based Antibody-drug Conjugate Targeting AXL-expressing Cancers. Mol Cancer Ther. 2022; 21(4):582–593.

51. Barretina J, Caponigro G, Stransky N, et al. The Cancer Cell Line Encyclopedia enables predictive modelling of anticancer drug sensitivity. Nature. 2012; 483(7391):603-607.

52. Wolpaw AJ, Grossmann LD, Dessau JL, et al. Epigenetic state determines inflammatory sensing in neuroblastoma. Proc Natl Acad Sci U S A. 2022; 119(6).

53. Sengupta S, Das S, Crespo AC, et al. Mesenchymal and adrenergic cell lineage states in neuroblastoma possess distinct immunogenic phenotypes. Nat Cancer. 2022; 3(10):1228–1246.

54. Borriello L, Nakata R, Sheard MA, et al. Cancer-Associated Fibroblasts Share Characteristics and Protumorigenic Activity with Mesenchymal Stromal Cells. Cancer Res. 2017; 77(18):5142–5157.

55. Hwang TJ, Orenstein L, DuBois SG, Janeway KA, Bourgeois FT. Pediatric Trials for Cancer Therapies With Targets Potentially Relevant to Pediatric Cancers. J Natl Cancer Inst. 2020; 112(3):224–228.

56. Zhang Z, Lee JC, Lin L, et al. Activation of the AXL kinase causes resistance to EGFR-targeted therapy in lung cancer. Nat Genet. 2012; 44(8):852–860.

57. Debruyne DN, Bhatnagar N, Sharma B, et al. ALK inhibitor resistance in ALK(F1174L)-driven neuroblastoma is associated with AXL activation and induction of EMT. Oncogene. 2016; 35(28):3681–3691.

58. Byers LA, Diao L, Wang J, et al. An epithelial-mesenchymal transition gene signature predicts resistance to EGFR and PI3K inhibitors and identifies Axl as a therapeutic target for overcoming EGFR inhibitor resistance. Clin Cancer Res. 2013; 19(1):279–290.

59. Eleveld TF, Oldridge DA, Bernard V, et al. Relapsed neuroblastomas show frequent RAS-MAPK pathway mutations. Nat Genet. 2015; 47(8):864–871.

60. Schramm A, Koster J, Assenov Y, et al. Mutational dynamics between primary and relapse neuroblastomas. Nat Genet. 2015; 47(8):872–877.

61. Gartlgruber M, Sharma AK, Quintero A, et al. Super enhancers define regulatory subtypes and cell identity in neuroblastoma. Nat Cancer. 2021; 2(1):114–128.

62. Yarmarkovich M, Marshall QF, Warrington JM, et al. Targeting of intracellular oncoproteins with peptide-centric CARs. Nature. 2023; 623(7988):820-827.

63. Liu X, Song J, Zhang H, et al. Immune checkpoint HLA-E:CD94-NKG2A mediates evasion of circulating tumor cells from NK cell surveillance. Cancer Cell. 2023; 41(2):272–287 e279.

64. Pereira BI, Devine OP, Vukmanovic-Stejic M, et al. Senescent cells evade immune clearance via HLA-E-mediated NK and CD8(+) T cell inhibition. Nat Commun. 2019; 10(1):2387.

65. Ravindranath MH, Filippone EJ, Devarajan A, Asgharzadeh S. Enhancing Natural Killer and CD8(+) T Cell-Mediated Anticancer Cytotoxicity and Proliferation of CD8(+) T Cells with HLA-E Monospecific Monoclonal Antibodies. Monoclon Antib Immunodiagn Immunother. 2019; 38(2):38–59.

66. Cascone T, Kar G, Spicer JD, et al. Neoadjuvant Durvalumab Alone or Combined with Novel Immuno-Oncology Agents in Resectable Lung Cancer: The Phase II NeoCOAST Platform Trial. Cancer Discov. 2023; 13(11):2394–2411.

67. Paraiso KH, Das Thakur M, Fang B, et al. Ligand-independent EPHA2 signaling drives the adoption of a targeted therapy-mediated metastatic melanoma phenotype. Cancer Discov. 2015; 5(3):264–273.

68. Noronha A, Belugali Nataraj N, Lee JS, et al. AXL and Error-Prone DNA Replication Confer Drug Resistance and Offer Strategies to Treat EGFR-Mutant Lung Cancer. Cancer Discov. 2022; 12(11):2666–2683.

69. Corbacioglu S, Steinbach D, Lode HN, et al. The RIST design: A molecularly targeted multimodal approach for the treatment of patients with relapsed and refractory neuroblastoma. Journal of Clinical Oncology. 2013; 31(15_suppl):10017–10017.

70. Koopman LA, Terp MG, Zom GG, et al. Enapotamab vedotin, an AXL-specific antibody-drug conjugate, shows preclinical antitumor activity in non-small cell lung cancer. JCI Insight. 2019; 4(21).

71. Van Renterghem B, Wozniak A, Castro PG, et al. Enapotamab Vedotin, an AXL-Specific Antibody-Drug Conjugate, Demonstrates Antitumor Efficacy in Patient-Derived Xenograft Models of Soft Tissue Sarcoma. Int J Mol Sci. 2022; 23(14).

72. Boshuizen J, Koopman LA, Krijgsman O, et al. Cooperative targeting of melanoma heterogeneity with an AXL antibody-drug conjugate and BRAF/MEK inhibitors. Nat Med. 2018; 24(2):203–212.

73. Boshuizen J, Pencheva N, Krijgsman O, et al. Cooperative Targeting of Immunotherapy-Resistant Melanoma and Lung Cancer by an AXL-Targeting Antibody-Drug Conjugate and Immune Checkpoint Blockade. Cancer Res. 2021; 81(7):1775–1787.

